# A Translaminar Spacetime Code Supports Touch-Evoked Traveling Waves

**DOI:** 10.1101/2024.05.09.593381

**Authors:** Daniel L. Gonzales, Hammad F. Khan, Hayagreev V.S. Keri, Saumitra Yadav, Christopher Steward, Lyle E. Muller, Scott R. Pluta, Krishna Jayant

## Abstract

Linking sensory-evoked traveling waves to underlying circuit patterns is critical to understanding the neural basis of sensory perception. To form this link, we performed simultaneous electrophysiology and two-photon calcium imaging through transparent NeuroGrids and mapped touch-evoked cortical traveling waves and their underlying microcircuit dynamics. In awake mice, both passive and active whisker touch elicited traveling waves within and across barrels, with a fast early component followed by a variable late wave that lasted hundreds of milliseconds post-stimulus. Strikingly, late-wave dynamics were modulated by stimulus value and correlated with task performance. Mechanistically, the late wave component was i) modulated by motor feedback, ii) complemented by a sparse ensemble pattern across layer 2/3, which a balanced-state network model reconciled via inhibitory stabilization, and iii) aligned to regenerative Layer-5 apical dendritic Ca^2+^ events. Our results reveal a translaminar spacetime pattern organized by cortical feedback in the sensory cortex that supports touch-evoked traveling waves.

**GRAPHICAL ABSTRACT AND HIGHLIGHTS:** 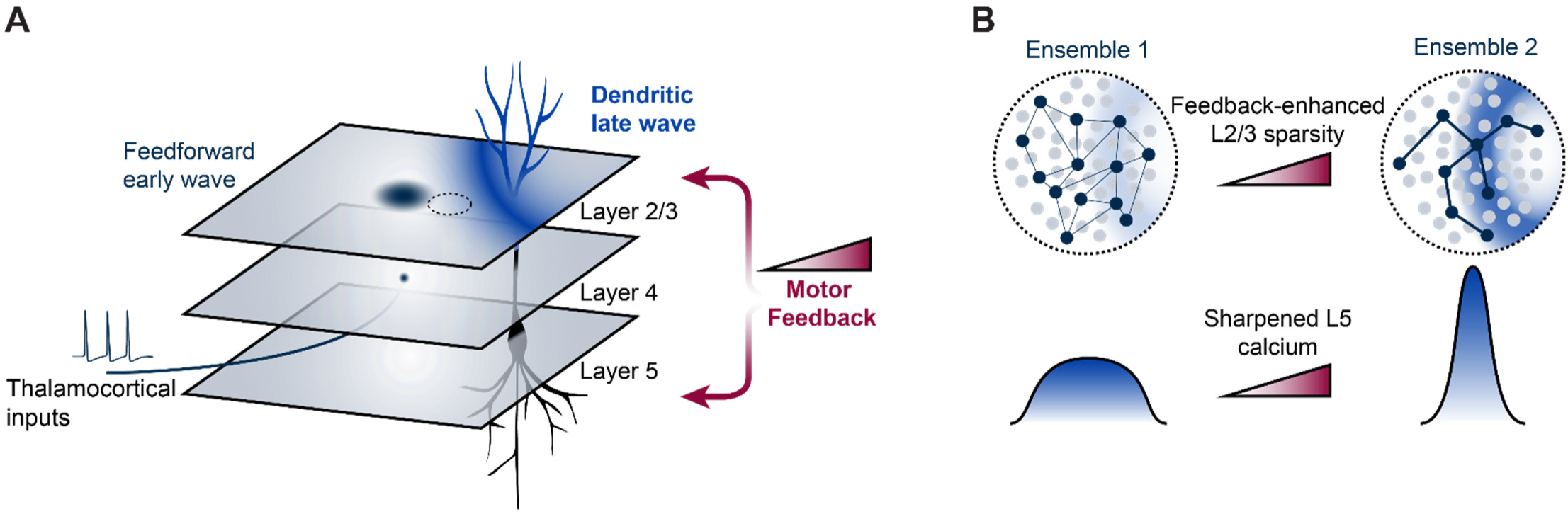

- Whisker touch evokes both early- and late-traveling waves in the barrel cortex over 100’s of milliseconds
- Reward reinforcement modulates wave dynamics
- Late wave emergence coincides with network sparsity in L23 and time-locked L5 dendritic Ca^2+^ spikes
- Experimental and computational results link motor feedback to distinct translaminar spacetime patterns

## INTRODUCTION

Traveling waves are a fundamental and widespread feature of neuronal activity in mammalian brains^1–9^. These patterns are an emergent property of recurrently connected systems^6,10,11^ and are believed to play a crucial role in modulating the excitability patterns of cells through precise phase delays, thus coordinating local and global neural activity^4^. Notably, both evoked and spontaneous traveling waves have been implicated in various cognitive functions such as sensory processing^1,10,12^, motor control^13^, attention, and working memory^3^. Recent evidence suggests that spontaneous cortical activity organizes in the form of traveling waves, shapes stimulus-evoked responses and predicts behavioral output^5^. In each of these cases, the circuit patterns accompanying and supporting traveling waves remain unclear^7,14^, an aspect we address in this study.

There are currently two main theories for the neural code underlying sensory-evoked cortical computations: the temporal code and the rate code^15^. Both of these theories, however, leave open the precise definition of the “window” or “interval of time” over which behaviorally relevant sensory stimuli are encoded in the cortex. By structuring neural activity in single cortical regions across space and time, traveling waves may serve as a substrate to help bind and integrate information over the biologically important periods when sensory-driven computations occur, leading to a spacetime code. Specifically, this spacetime code refers to a situation in which the spike rates and times of a recurrent neural population at any given moment represent sensory-evoked activity that occurred over an interval of time, enabled by waves.

How do neural circuits support such waves? Computational evidence suggests that axonal fiber delays due to long-range axons traversing superficial layers in the cortex promote traveling waves^4^. However, how these traveling waves shape translaminar circuit dynamics and, in turn, how circuit patterns support their propagation remains unknown. Unraveling the link between sensory-evoked traveling wave dynamics and local microcircuit activity is crucial in comprehending how distributed neural computations encode sensory features and shape cognition.

A well-established model for studying sensory processing and related cognitive functions uses the neural response to whisker touch in mice^12,16^. This system is well suited to unmask the neural circuits underlying goal-directed behaviors. Previous studies across the primary whisker somatosensory cortex (wS1) in mice have shown that whisker touch elicits a radially propagating pattern of neural activity that spreads out from the point of maximal input in the cortex ^10,12^. Whether this barrel cortex activity is organized as a wave has never been experimentally tracked. A single-whisker deflection evokes a sensory response, driven by direct thalamic input, that initiates across multiple layers at its somatotopically aligned barrel in wS1^17^. This response rapidly spreads across adjacent barrels^18^, the secondary whisker somatosensory cortex (wS2), and the whisker motor cortex (wMC)^12^, including primary and secondary areas. Thus, the whisker barrel represents a powerful system to tease apart the role of short and long-range connectivity and translaminar circuitry to potential traveling wave spread. On the scale of single cortical regions, experimental evidence suggests that evoked waves allow neural circuits to flexibly store and use information from the recent past, a feature critical to integrating higher-level stimulus features distributed more broadly across space and time^7^. However, the precise pattern of circuits underlying and supporting touch-evoked wave activity over hundreds of milliseconds remains unknown.

Based on data from the barrel cortex, computational analysis suggests that a sparse network of strongly connected excitatory neurons in cortical layer 2/3 (L2/3) drives the emergence of sensory-evoked cortical waves^10,12,19^. This model implies that only a small subset of L2/3 neurons is necessary for the propagation of these waves despite the spatially intermixed response of this subpopulation. Other studies have furthered this model and suggested that these waves propagate through topographically organized, recurrent functional connections in horizontal unmyelinated axons^4^. The sparse wave here is defined as a network in which wave propagation is supported by just a few local cells that spike as the wave passes through. Even though network sparsity in L2/3 is considered a hallmark of traveling waves, observing this “sparse wave” phenomenon directly using conventional recording technologies has been challenging. To do so, one must record superficial traveling waves while simultaneously mapping L2/3 cellular dynamics with a high spatial (cellular and subcellular) resolution.

Previous experiments on traveling waves relied on voltage dye imaging^14,20^, but this technique is currently limited to the superficial cortical layers and offers insufficient spatial resolution. While two-photon imaging allows cellular-scale imaging, it cannot capture the temporal dynamics of the wave. Some studies have used Utah Arrays^5^, macroscale ECoG arrays^21^, or microscale ECoG grids^22,23^ to capture traveling waves, but haven’t integrated such recordings with high-resolution functional imaging, limiting the ability to identify the sparse cortical ensembles and translaminar circuits underlying wave structure. In this study, we overcome this challenge in the mouse barrel cortex with a novel multimodal platform that provides high spatiotemporal resolution wave recordings and cellular-scale two-photon imaging.

We designed, fabricated, and customized flexible and transparent polymer-based NeuroGrids^22,24^ that allow recording traveling waves with high spatial resolution across multiple barrels. By combining this transparent µECoG array with simultaneous two-photon imaging, high-density silicon probe recordings, optogenetics, and pharmacology, we elucidated, for the first time, the translaminar cellular patterns supporting the traveling wave. We directly observed changes in L2/3 sparsity during different traveling wave dynamics. We identified a late, sensory-evoked traveling wave that requires wMC ^25^ and coincides with calcium plateaus in the apical dendrite of L5 neurons^26–28^. We reconcile this motor cortical effect through a recurrent balanced-state spiking network model that incorporates realistic neuronal densities, connection probabilities, axonal delays, and synaptic conductances, to achieve an inhibitory stabilized network. Taken together, this study establishes a link between motor cortical feedback, L2/3 network sparsity and the activation of L5 apical dendrites as key elements underlying stimulus-evoked traveling waves.

## RESULTS

### NeuroGrids for mapping traveling wave dynamics during two-photon imaging

We used scalable semiconductor manufacturing to create flexible and semi-transparent NeuroGrids^22,23,29^ (**Figure 1A-B**, **see Methods**). These parylene-based^30^ µECoG interfaces were patterned with an array of cellular-scale recording sites (25 µm diameter) coated with a specialized layer of nanoporous gold^31,32^ for low-impedance recording (**Figure 1B-C, Figure S1A-C, see Methods**). Although the NeuroGrids can be customized in several ways such as the length, size, electrode density, and through-hole access, we designed them specifically for high-resolution recordings across 2-4 barrels in mouse barrel cortex.

**Figure 1.**
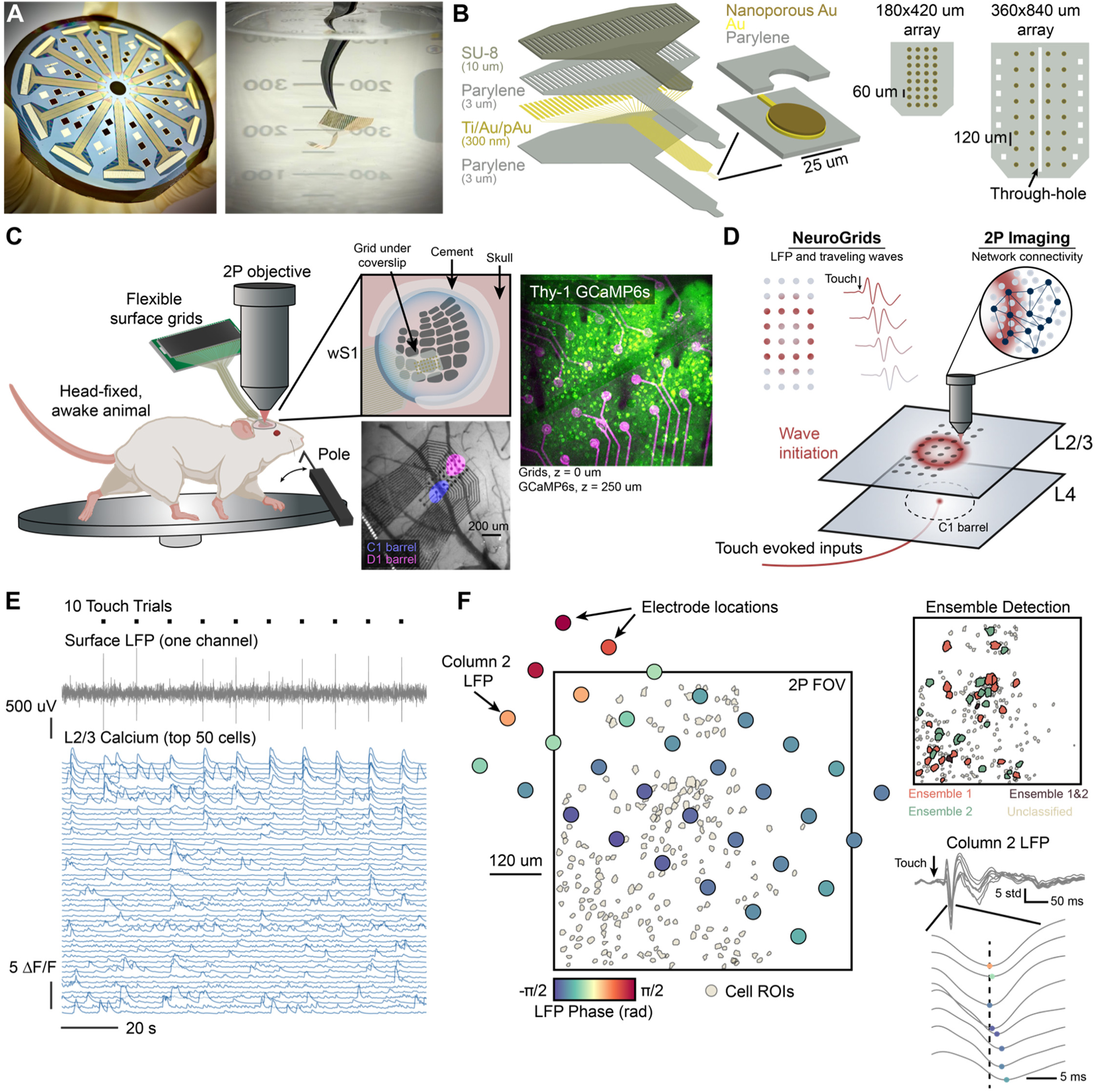
Flexible, transparent grids for simultaneous traveling wave detection and two-photon imaging – An overview. **(A)** (Left) 4” silicon wafer with 12 surface grids fabricated simultaneously. After fabrication, the flexible grids are peeled from the silicon substrate. (Right) Grid held in water to demonstrate flexibility. **(B)** (Left) Cross-sectional stack of the fabricated surface grids. A thin-film parylene substrate is followed by a SU-8 thickening layer to stiffen the back end of the probe to prevent excessive bending and folding. Thin-film Au forms the metal leads, and an extra 400 nm of nanoporous gold is deposited onto the recording sites for reduced electrochemical impedance. (Right) To-scale schematics of the surface grid designs used in this work. **(C)** Schematic depicting simultaneous surface electrophysiology and two-photon imaging. Awake, head-fixed mice run on a wheel and receive either passive or active whisker touch. Surface grids are acutely implanted over wS1. Top inset, shows that we place surface grids onto the brain and seal the cranial window with a glass coverslip. The bottom inset is an image of grids on the brain overlaid with the results of intrinsic-optical imaging used to locate the C1 and D1 barrels. Right inset, validation of the transparent nature of two-photon imaging through the grids. **(D)** Schematic representing the different recording schemes. Touch evokes traveling waves of field potentials in the upper cortical layers are mapped with NeuroGrids. Concurrently, we also capture cellular-scale network activity with two-photon imaging. **(E)** Example data showing 10 consecutive passive whisker-touch trials, LFP recordings from one channel on the grid array, and calcium traces from 50 select cells in the field of view. **(F)** (Left) To-scale schematic showing the exact NeuroGrid locations overlaid with the two-photon field of view and detected ROIs 200 µm below the cortical surface. Electrode colormap depicts the average phase of the LFP in a short 10 ms window following touch. (Right, top) shows the same FOV, but the cell ROIs are color-coded to match identified ensembles detected with functional connectivity analysis (see Methods). Ensemble ROIs are increased in size by 200% for clarity. (Right, bottom) a representative example showing the average LFP across column 2 of the electrode array depicting a progressive traveling wave.

The transparency of our NeuroGrids allowed us to combine surface recordings with simultaneous multiphoton imaging (**Figure 1C-F**). After locating the barrel target with intrinsic optical imaging^33^, we placed NeuroGrids on the cortical surface and sealed across the cranial window with a conventional coverslip (**Figure 1C**, **see Methods**). This process along with the porous properties of our electrodes allowed us to perform L2/3 two-photon calcium imaging directly through the grids in awake, head-fixed mice (**Figure 1D-E**) with minimal impacts on imaging quality^34^. Noise artifacts from laser scanning were minimal when imaging GCaMP6s, and were easily filtered out with a recursive notch filter^35^ (**Figure S1D-E**). While alternative techniques can be used to further reduce noise artifacts, such as using red-shifted GCaMP sensors, or enhancing the electrode transparency with graphene^29,36^; nanomesh arrays^37^, or other novel materials^38–40^, our recordings revealed that nanoporous gold was sufficient for near artifact-free imaging while allowing for dense patterning. This platform allowed us to record from hundreds of L2/3 cells in the field of view (FOV) with clearly detectable touch-evoked LFP (**Figure 1E**, **Figure S1F-G**).

Whisker touch (**see Methods**) evoked strong LFP and calcium transients throughout the barrel and surrounding areas (**Figure 1E-F**). The LFP was non-stationary and showed clear latencies across the NeuroGrid, indicative of traveling waves (**Figure 1F**). By recording simultaneous LFP and cellular calcium in dozens of trials, we could construct maps of LFP latency across the grid in combination with maps of consistent L2/3 network ensembles that arose throughout the recordings (**Figure 1F, see Methods**). This powerful multimodal platform allowed us to dissect the underlying circuit patterns during cortical traveling waves, described below under passive and active touch.

### Passive touch evokes early and late waves in barrel cortex

We first teased apart touch-evoked traveling wave dynamics from an electrophysiological perspective using NeuroGrids (**Figure 2A**). In awake animals, passive C1 whisker deflection (**see Methods**) drove clear patterns of wave-like of activity across the principal barrel and surrounding columns (**Figure 2A, bottom**). Here, passive touch was presented in the absence of reward to avoid confounding effects of anticipation^41^ and history-based reward accumulation^42^ on wave dynamics, which can impact cortical responses in a history-dependent manner^7,43^. Averaging LFP across trials can distort and even obscure traveling wave activity^7^. Therefore, to detect and quantify these waves on a single-trial basis, we used an approach for wave detection called “generalized phase” (GP, **Figure 2B, Figure S2A-C, see Methods**) ^5^. Like conventional analyses that use a Hilbert transform to attain the analytic signal and instantaneous phase, the GP allows obtaining instantaneous phase of a wideband signal (3-40 Hz, see Methods) by addressing technical limitations in this signal processing framework. This approach is advantageous, as it can quantify phase (which can, in turn, be used to detect traveling waves) without requiring the assumption that the LFP is narrowband (and, in turn, potentially introducing filtering artifacts). When applied to NeuroGrid recordings, our GP pipeline estimated the instantaneous phase of the wideband LFP and detected both spontaneous and touch-evoked waves (**Figure 2C**, **see Methods**). Across all animals (n = 9), we observed spontaneous activity, robustly stereotyped early waves within 50 ms of touch, and a burst of delayed waves (**Figure 2D**). Strikingly, within the first ∼100 ms post touch, we observed nested epochs with periods comprising of waves and no waves, signifying intermingled periods of zero-lag phase (perfectly synchronized) and desynchronized (un-organized phase / with phase delays) LFPs across the grids, respectively. Importantly, though, post ∼100 ms, we observed a wave pattern emerge across the surface with a distinct spatiotemporal structure. This delayed potential initiated ∼100 ms after touch and peaked ∼150- 200 ms after touch (**Figure 2E**). We termed these delayed dynamics “late waves” and the patterns were distinctly different from the early components in directionality (**Figure 2F**). It is critical to note that this late wave persisted much after the stimulus presentation, suggesting reverberatory effects across recurrent networks. We also tested the possibility that changes in locomotion could be attributed to the wave potentials. Still, we found no strong correlation between the surface LFP and wave times to motion initiation (**Figure S2D-E**). Importantly, our mice were stationary and not locomoting during passive touch trials (**Figure S2F**) and we also found a significant reduction in surface potential and wave dynamics when applied to periods in which the mouse freely whisked, i.e. whisker motion onset evoked (not shown). We quantified speeds in the pre-stimulus period, early wave period, and during the late wave and found that the early waves traveled slightly faster in comparison to the late wave (**Figure 2G**, p < 0.001, n = 9 animals, 289 spontaneous waves, 486 early waves, 411 late waves, Kruskal-Wallis with a *post-hoc* Dunn-Sidak test). Recent works have alluded to the idea of inhibition in reducing traveling wave speed^44^, and others have suggested inhibition is critical for timing^45^. This would allude to the possibility that the late wave dynamic could be associated with inhibitory restraint on the cortical patch, leading to overall spatio-temporal synchronization across layers.

**Figure 2.**
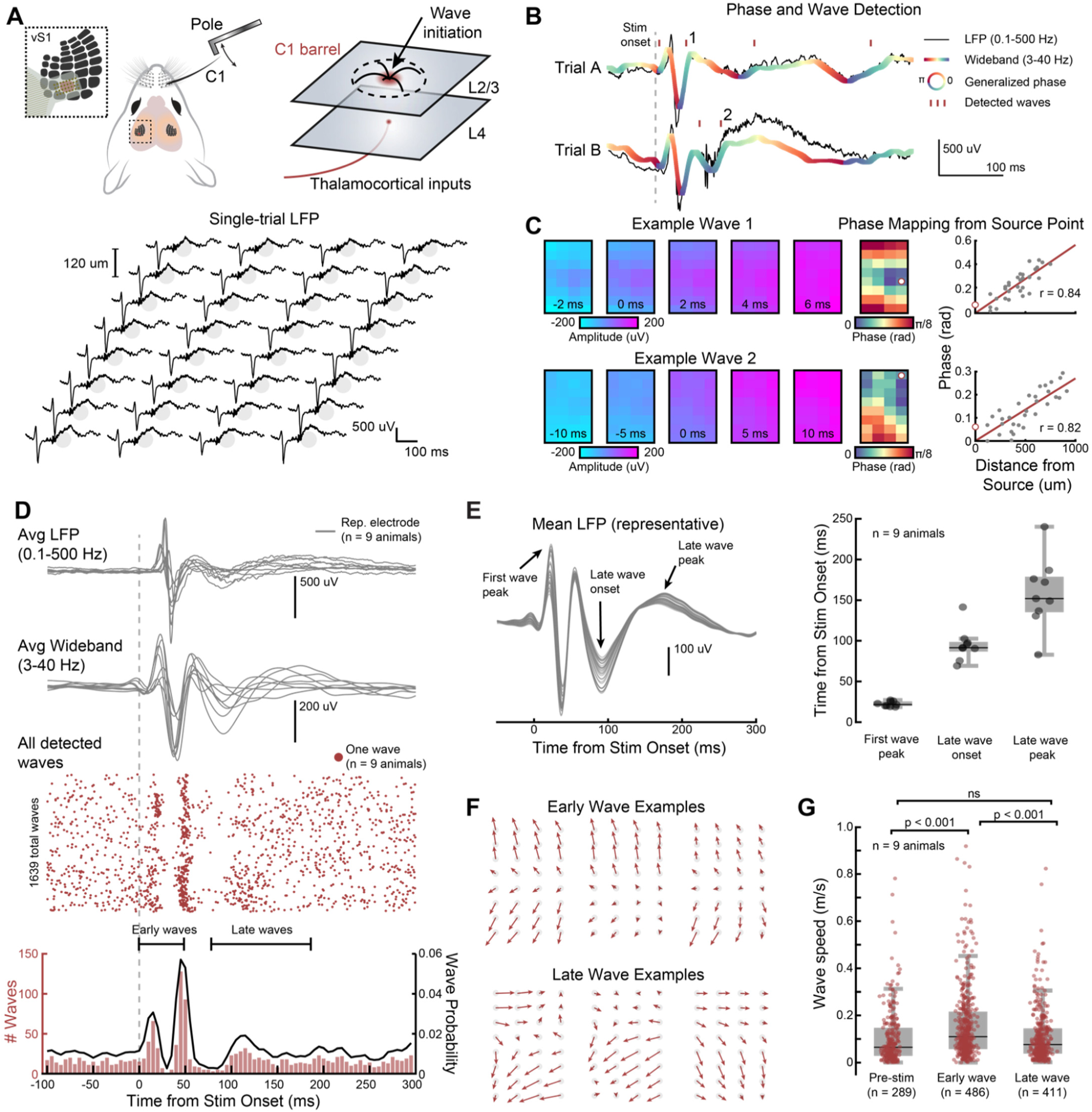
Whisker touch evokes early and late traveling waves across the barrel cortex. **(A)** (Top) Schematic of the experimental setup. We used intrinsic imaging to target the C1 barrel with NeuroGrids and delivered passive whisker stimulation to the C1 whisker. (Bottom) Evoked LFP across the NeuroGrid from a single touch stimulation. **(B)** Two representative touch trials detected traveling waves using the generalized phase (GP) approach. LFP is from a representative channel on the NeuroGrid. The gray line indicates the full LFP spectrum (0.1-500 Hz). The thick color line is the wideband LFP signal, filtered from 3-40 Hz for traveling wave detection and analysis. Colormap is the calculated generalized phase (GP) of the wideband signal. Red dashes indicate detected waves (see Methods). **(C)** (Left) Example waves 1 and 2 indicated in (B) traveling across the NeuroGrid. (Right) For each wave, we compute a phase map and wave source point (white dot, see Methods). These act as a snapshot of wave propagation. For detected waves, the phase across the grid strongly correlates with distance from the source point (see also Figure S2C). **(D)** Average LFP, wideband LFP, and detected traveling waves around the stimulus period for all animals following whisker touch. Only the LFP from one representative channel is shown for each animal. LFP is aligned to the onset of pole motion (n = 9 animals, 1639 total detected waves across 607 trials). We observe strong “early” sensory-evoked traveling waves <50 ms after the stimulus, in addition to a second bump of “late” traveling waves ∼100-150 ms after touch. **(E)** (Left) Mean grid LFP (3-40 Hz) across all trials for a representative animal with distinct features noted. (Right) Quantified times for each feature in the average LFP signal using a representative electrode (n = 9 animals). **(F)** Vector plots showing wave propagation in a representative animal. Note that the early waves are stereotyped, and the late waves are more variable. **(G)** Traveling wave speed for detected waves detected in the pre-stimulus, early-evoked, and late-evoked periods (n = 9 animals, 289 pre-stimulus waves, 486 early waves, 411 late waves; ns = not significant, p < 0.001 Kruskal-Wallis with a post-hoc Dunn-Sidak test). Values are in good agreement with propagation speeds along unmyelinated axons.

### Rewarding action in a goal-directed processing task modulates early and late traveling wave dynamics

Our results with passive-touch stimulation led us to probe whether traveling waves across the barrel cortex can reflect behaviorally relevant information under active paradigms where volitional movement of whiskers against an object occurs. We hypothesized that reward-based reinforcement would modulate wave properties towards dynamics that enhance the saliency of a reward stimulus under active touch. To test this, we developed a two-whisker, active-touch discrimination task (**Figure 3A-C**, **Figure S3A, see Methods**). In head-fixed animals, we targeted the C1 and D1 whiskers separately with a system of pneumatic pistons^46^ that animals were free to actively whisk against for 1.3 s (**Figure 3B-D**, **Figure S3A, see Methods**). We trained animals in an operant task to lick for a water reward during C1 touch (Go stimulus) and withhold licks during D1 touch (No-Go stimulus, **Figure 3C**, **Figure S3A**). In this scenario, Go and No-Go trials should evoke waves of activity, including actively engaging higher-order feedback in the C1 and D1 barrels, allowing us to test for goal-directed changes to traveling waves (**Figure 3A**). This is further strengthened by the growing evidence that higher-order wMC feedback plays a role in shaping behavior^47^ and is fundamental during active-touch^48^

**Figure 3.**
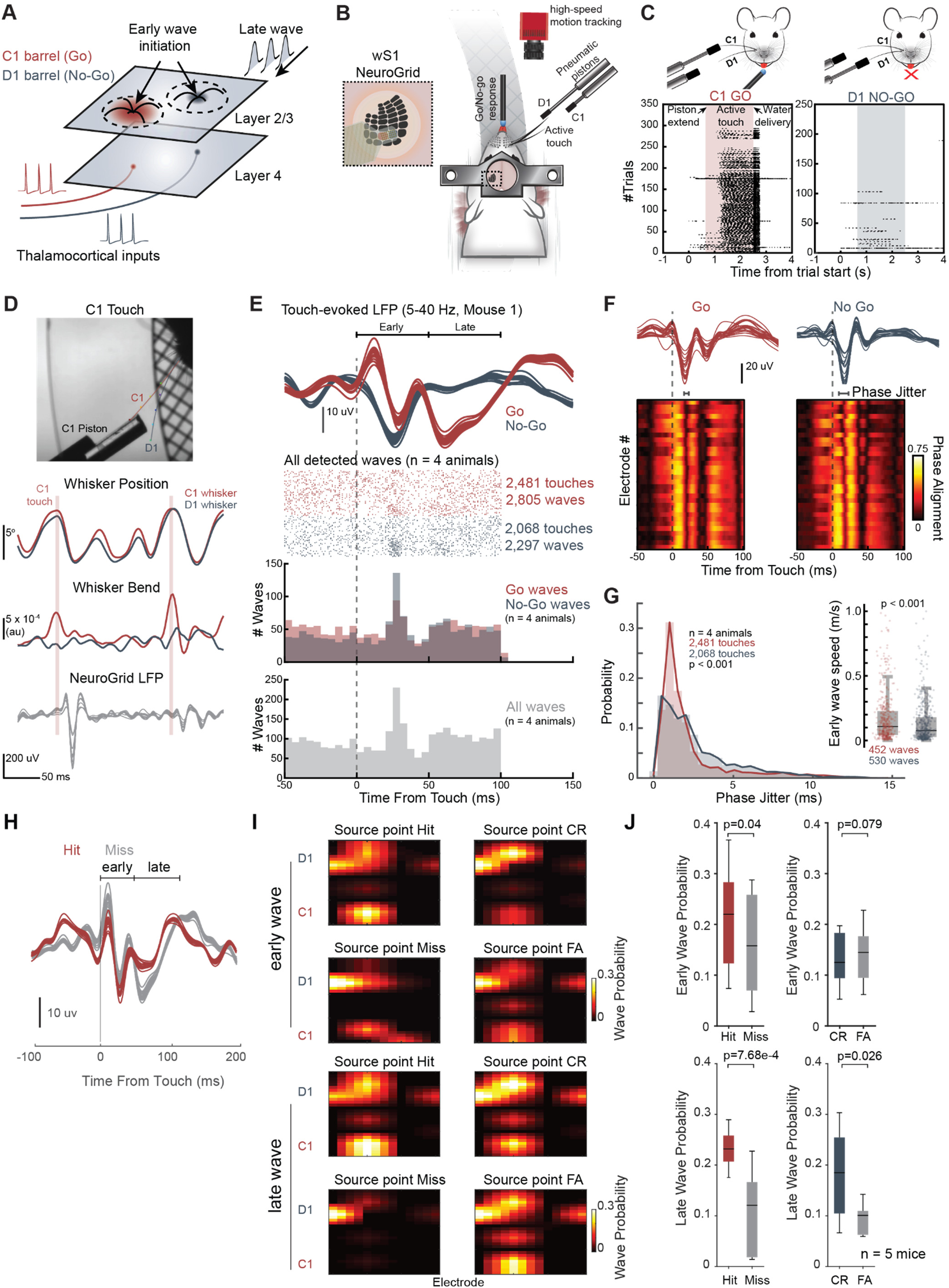
Rewarding action in a Go-NO-Go paradigm modulates early and late traveling wave dynamics during active touch. **(A)** Schematic depicting traveling wave initiation and late wave reverberation in two adjacent barrels, C1 and D1, corresponding to Go and No-Go whisker stimuli, respectively. **(B)** (Left) Schematic of our two-whisker active-touch paradigm. Head-fixed animals run on a wheel while pneumatic pistons extend into the whisking range of the C1 (Go stimulus) or D1 (No-Go stimulus) whiskers. Animals have been trained to lick for a water reward during the Go stimulus. We recorded whisker kinematics with high-speed videography and wS1 dynamics with NeuroGrids (inset) during this experiment. **(C)** Lick raster plot for a representative animal during Go and No-Go trials. The shaded area indicates when each piston is extended into the whisking field. Each black dot is a single lick. **(D)** Representative whisker kinematics with simultaneous NeuroGrid recordings. Top micrograph shows a DeepLabCut pose estimate of the C1 and D1 whiskers during C1 touch. This tracking quantifies whisker kinematics such as position (top trace) and whisker bending (middle trace), and allows for detecting precise single-whisker touch times. We use these touch times to analyze the touch-evoked LFP (bottom trace). **(E)** We applied our traveling wave detection pipeline to the touch-evoked LFP during C1 and D1 touch. We only analyzed touch times with a separation of more than 100 ms so that rapid consecutive touches do not interfere with our analysis during the late wave period. (Top) Mean touch-evoked LFP (5-40 Hz) for all grid electrodes for a representative animal. Red are Go trials, and gray are No-Go trials. Note the significant late-wave LFP during Go trials. (Middle-Top) Raster plot of all detected traveling waves across animals during Go and No-Go trials. (Middle, bottom) Histogram of all detected waves across animals during Go and No-Go trials. (Bottom) Histogram of all detected waves for all trials combined. (n = 4 animals, 2,481 Go touches, 2,805 Go waves, 2,068 No-Go touches, 2,297 No-Go waves). **(F)** (Top) Mean touch-evoked LFP (5-40 Hz) for all grid electrodes for a representative animal. (Bottom) Mean phase alignment for each electrode across the NeuroGrid (see Methods). In this analysis, periods of high alignment with structured delays are associated with traveling waves. Both Go and No-Go touch-evoked LFP have intermittent periods of high phase alignment. An increase in jitter across the grid for the No-Go condition based on phase alignment is observed. **(G)** Quantified touch-evoked phase jitter (n = 4 animals, 2,481 Go touches, 2,068 No-Go touches, p < 0.001 using a ranked-sum Wilcoxon test following a bootstrap with 1000 iterations for each animal (see Methods). (Inset) The shift in phase jitter for Go trials is reflected by increased wave speed compared to No-Go trials (n = 4 animals, 452 Go waves, 530 No-Go waves, p < 0.001, ranked-sum Wilcoxon test). **(H)** Surface LFP response during hit-and-miss response across the grid for the Go whisker (C1). **(I)** Probability of wave source points for a grid covering both barrels. Notice how probability is somatotopically mapped depending on Go (C1) vs No Go (D1) trials. We also observed a clear difference in probability as a function of task outcome during the late wave period, specifically during the miss. **(J)** (top) Early traveling wave probability for each task outcome (n = 4 animals, 2,481 Go touches, 2,068 No-Go touches). (bottom) Late traveling wave probability for each task outcome. (n = 4 animals, 452 Go (hit) waves, 530 No-Go (CR) waves, ranked-sum Wilcoxon test)

After training animals to expertly discriminate (**Figure S3B**), we performed NeuroGrid surface recordings and high-speed whisker imaging as animals undertook the behavioral task. We then used whisker tracking with DeepLabCut^49^ to detect the precise C1 and D1 touch times during Go and No-Go trials (**Figure 3D, see Methods**). As with passive-touch recordings, we used our GP pipeline to detect spontaneous and touch-evoked traveling waves in the wideband filtered LFP throughout the experiment. We restricted our analysis to consecutive touches with an interval greater than 100 ms to remove the confounding factor of multiple touches happening in rapid succession, which we believed would interfere with late wave detection. In results not shown, however, we found that only the first touch resulted in an appreciable surface potential swing, while consecutive touches (4-8Hz) did not create additional large depolarizations. Instead, it appeared that the large surface potential swing and wave dynamic created by the first touch opened a 50 to 200 ms window over which the cortical brain state was modified for consecutive touches.

Mirroring our results with passive touch, across all animals we detected both early (< 50 ms after touch) and late (50-100 ms after touch) touch-evoked traveling waves in both Go and No-Go trials (**Figure 3E**, n = 4 animals, 2,481 C1 touches, 2,068 D1 touches, 2,805 Go waves, 2,297 No-Go waves). We confirmed these waves results are touch-related by analyzing waves during free whisking in the absence of touch and did not observe a strong correlation between LFP potentials or traveling wave onset times and whisking phase (**Figure S3C-D**, n = 5 animals, 5507 total whisking cycles, 537 total detected waves). We also found that touch-evoked waves exhibited significantly larger LFP amplitudes (**Figure S3D**, n = 5 animals, 537 whisking-only waves, 1275 early waves, 963 late waves). We did observe that the late wave analysis window (**Figure 3E**, 50-100ms) was at odds with the results in Figure 2D (passive touch), where 50-100ms is the period with the least number of waves detected before persistent wave activity was observed. Given the trough-to-peak in the surface potential signaled the onset of waves, we surmised that the LFP and wave structure during active touch is more compressed as the trough-to-peak occurs earlier, possibly due to differences in brain state and local circuit dynamics.

How does goal-directed behavior shape touch-evoked phase dynamics within the barrel? We quantified specific properties of phase across the NeuroGrid during Go and No-Go trials to determine the effects of sensory associations on traveling waves. We discovered a notable difference in phase jitter across the NeuroGrid, specifically a sharpened phase distribution during Go trials compared to No-Go trials (**Figure 3F-G**, n = 4 animals, 2,481 Go touches, 2,068 No-Go touches, p < 0.001, ranked-sum Wilcoxon test). Such synchronization effects were reflected in wave speed differences between Go and No-Go touches during the early (p < 0.001) and late period (p < 0.01, data not shown), with Go waves slightly faster than No-Go waves (**Figure 3G** inset, n = 4 animals, 452 early Go waves, 530 early No-Go waves, ranked-sum Wilcoxon test). We also noted a significant enhancement of the late LFP for the Go stimulus (**Figure S3E-F**) (n = 4 animals, p < 0.001, Wilcoxon signed-rank test). These results indicate that rewarding action drives distinct wave properties, likely influencing synchronization across the translaminar axis, and that the late wave may be a distinct mechanism for more accurate sensory perception, particularly because, whisker movement was largely stereotyped across both the Go and No-Go trials (**Figure S3G-J**).

Indeed, upon further analysis, and significantly, we found that traveling wave source points were distinctly localized to barrels, which encoded both hits, misses, and correct rejects (**Figure 3H, 3I**). A clear difference in the wave structure (timing, amplitude) and probability was observed for the hits vs. misses (**Figure 3H**), most noticeable during the late wave period across the principal whisker column (**Figure 3I, bottom**), an effect one would not observe by just measuring LFP alone. Moreover, the timescales of the wave and animal behavior suggest that the late wave may be a critical circuit feature for ensuring the correct initiation or withholding of a lick. The sensory-evoked waves occured over a ∼200ms window, and anticipatory licking ensued approximately 500ms after the first touch and lasted ∼1sec until reward delivery (**Figure 3C**). This timeline suggests that the late wave dynamic may not only be a distinct signature of salient stimuli, but also an important feature needed for optimal signal routing to ensure an impending lick or an intent to withhold response. To our knowledge, this is the first report showing cortical traveling wave structure modulation due to sensory associations. Overall, the wave property and not just the amplitude of the LFP—most notably during the late period—was strongly linked to future task outcome (**Figure 3J**). These experiments demonstrate that early and delayed sensory-evoked waves are apparent during active touch and carry context-dependent information critical for impending behavior.

### Touch-evoked waves across frequency bands reveal translaminar circuit contributions

Next, we focused on dissecting the underlying circuit dynamics that support a wave structure. We used single-whisker passive touch due to its tractability and simplicity for in-depth circuit interrogation. We analyzed specific frequency bands that are known to be associated with signal routing across specific cortical layers^50^. How does each frequency band support touch-evoked traveling waves observed on the NeuroGrid?

To reveal this, we re-analyzed traveling waves during the early and late periods, but with an emphasis on the narrowband frequencies involved (**Figure 4, Figure S4A**). Decomposing the full LFP spectrum into the gamma (30-90 Hz), beta (15-30 Hz), and theta (4-12 Hz) bands (see **Methods**) exposed a clear frequency dependence of the early and late waves, with an emphasis of the beta and theta bands on the late period (**Figure 4A**). When we detected traveling waves with our GP pipeline, we observed an intricate interleaving of touch-evoked wave events across frequency bands (**Figure 4B**, n = 9 animals, 14,598 gamma waves, 7,100 beta waves, 2,529 theta waves). Here, we observed that waves in the gamma band appeared to be nested within periods of beta waves. As with the wideband-filtered waves described in **Figure 2**, the median traveling wave speed of all detected narrowband waves falls within the reported range for unmyelinated axons (<1 m/s), with a progressive slowing in wave speeds in lower frequency bands (**Figure S4A**, n = 9 animals, 14,598 gamma waves, 7,100 beta waves, 2,529 theta waves).

**Figure 4.**
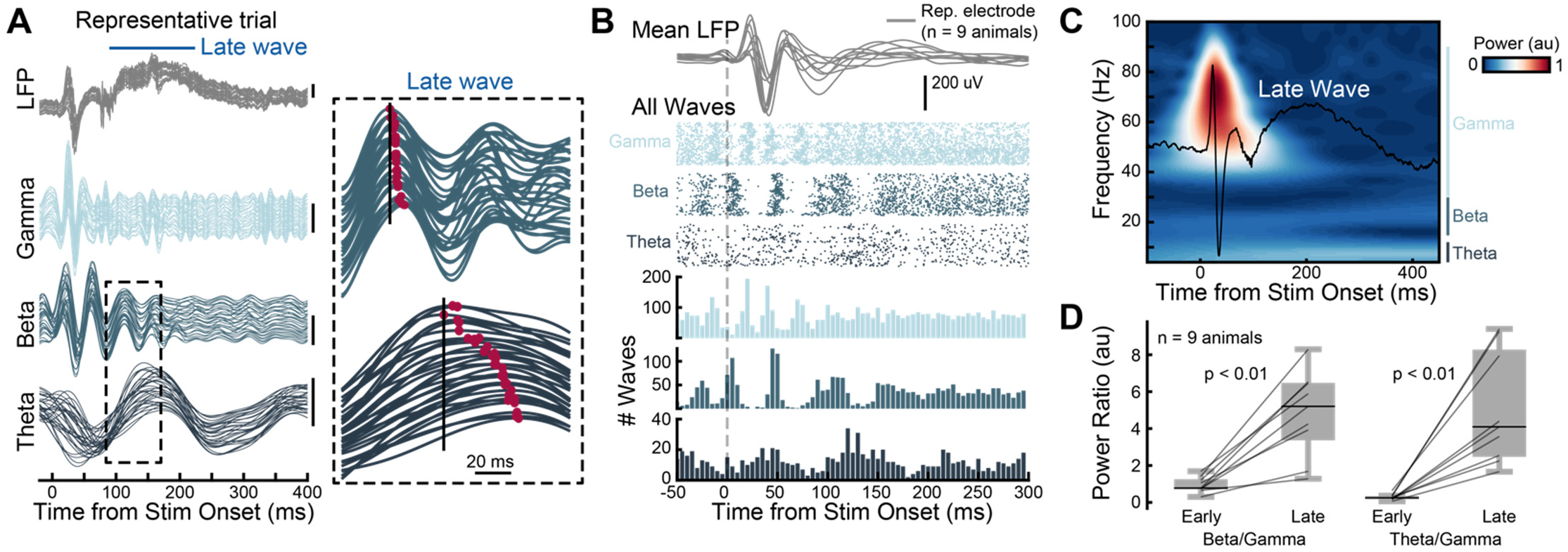
The beta and theta bands contribute to the late wave. **(A)** Single-trial LFP and the corresponding gamma (30-90 Hz), beta (15-30 Hz), and theta (4-12 Hz) frequency bands. The inset shows the beta and theta bands during the onset of the late wave. Red denotes the local maxima and shows wave latencies across the grid. **(B)** Detected traveling waves in the gamma, beta, and theta bands for all animals (n = 9 animals). Top traces show each animal’s mean touch-evoked LFP on the NeuroGrid on a representative electrode. The raster plot shows the onset times for all detected waves. The histogram shows the combined wave times across animals (n = 9 animals, 14,598 gamma waves, 7,100 beta waves, 2,529 theta waves). **(C)** Mean touch-evoked LFP on the NeuroGrid for a representative animal overlaid on the signal spectrogram. **(D)** Beta/Gamma and Theta/Gamma power ratios across animals in the early (0-50 ms) and late (100- 250 ms) windows following whisker touch (n = 9 animals, p < 0.01 signed-rank Wilcoxon test).

These interesting frequency-specific properties hint that the underlying circuits supporting sensory-evoked waves at the cortical surface are translaminar. Most conspicuously, with NeuroGrid recordings we found that the early-evoked waves (0-50 ms) are dominated by gamma-beta coupling, while the late wave (100-300 ms) is dominated by lower frequency beta-theta coupling (**Figure 4C-D**, n = 9 animals, p < 0.01, signed-rank Wilcoxon test). Previous studies in non-human primates have found a cortex-wide motif in which supra-granular circuits (L2/3) are dominated by the gamma band, and infragranular layers (L5/6) largely by beta and theta frequencies^50–52^. We replicated this finding in mice using silicon probe recordings alongside frequency analysis of touch-evoked LFP across cortical layers (**Figure S4B,** n = 3 animals). This analysis also revealed another important layer-specific motif. The late wave period (100-300 ms after touch) showed a marked increase in beta-theta power, but this low-frequency increase was largely restricted to the infragranular layers (**Figure S4C**). These combined observations provide strong evidence that the waves observed at the cortical surface involve an intricate translaminar coupling wherein infragranular circuits are important for the supragranular wave. Specifically, they point to distinct contributions across superficial and deep layer neurons, suggesting a precise spacetime code supports traveling wave dynamics and spread.

### Traveling waves and a L2/3 sparse code

We next sought to directly map layer-specific circuit dynamics underlying traveling waves, beginning with L2/3. Network sparsity in L2/3 has been implicated as a critical component underlying traveling wave propagation, albeit computationally^10^; however, existing experimental methods lack the multimodal capability necessary to directly observe this phenomenon. Here, we leveraged the transparency of the NeuroGrid array to combine our traveling-wave recordings with simultaneous two-photon imaging of L2/3 dynamics (**Figure 1C-F**, **Figure 5, see Methods**). Specifically, we hypothesized that different traveling wave dynamics—namely, trials with and without the prominent late wave—would be supported by different L2/3 circuits and possibly changes in network sparsity (**Figure 5A**). The presence and absence of a late wave occurred randomly, interleaved within trials in which there were appreciable late wave dynamics (52 ± 27% of all trials, 67 ± 21% of trials with a clearly detectable touch-evoked LFP, mean ± standard deviation, n = 13 animals). Waves were not linked to spontaneous whisking (**Figure S3C**) or behavioral state transitions (**Figure S2D-E**). Furthermore, since mice were stationary during passive whisker touch (**Figure S2F**), our results are not confounded by state-changes related to locomotion. We thus hypothesize that absence of late waves was most likely due to periodic brain state changes occurring on a trial-to-trial basis. Through the NeuroGrids, we consistently observed a strong touch-evoked LFP while imaging imaged hundreds of L2/3 cells at depths of 150- 250 um with a minimal impact on the number of cells in the FOV, calcium SNR, and number of touch-responsive L2/3 cells (**Figure 5B-C**, **Figure S1**).

**Figure 5.**
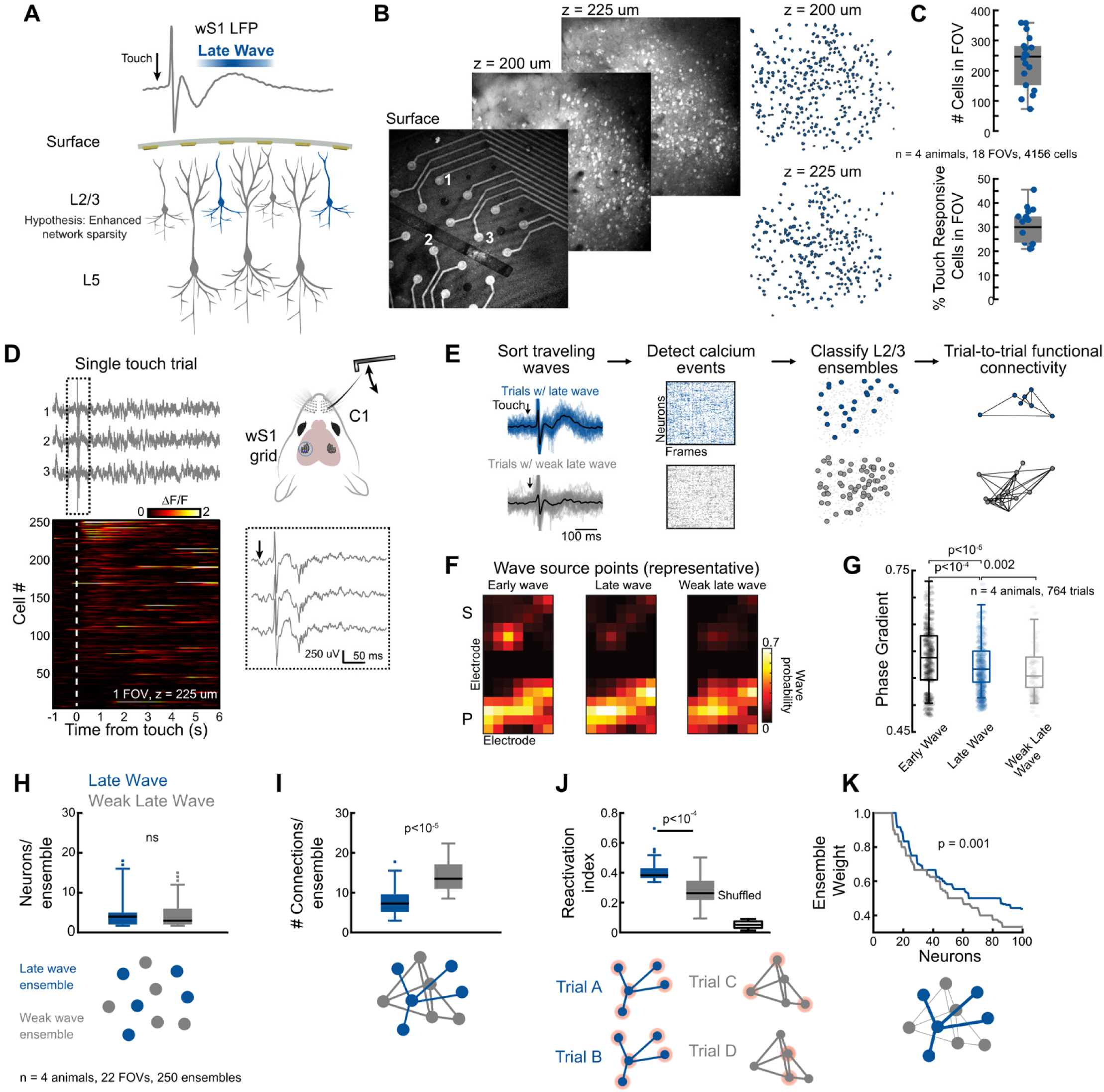
Enhanced L2/3 sparsity supports late wave dynamics. **(A)** Schematic of a sparse-network theory in L2/3 that supports the late wave. **(B)** Representative vS1 two-photon imaging through the surface grid. (Left) Images were taken at the cortical surface and depths 200 and 225 µm below the grids. (Right) Cell ROIs at each depth. **(C)** The number of cells in each field of view (FOV) across animals and the percent of touch-responsive cells in each FOV (see Methods, n = 4 animals, 18 total FOVs, 4156 total cell ROIs). **(D)** Representative C1 touch trial during simultaneous surface electrophysiology and 2P imaging at a depth of 225 µm below the grids. The electrophysiology traces correspond to the labeled electrodes in (B). The inset shows the electrophysiology traces immediately following touch. **(E)** 2P data analysis pipeline for detecting L2/3 neuronal ensembles and quantifying functional connectivity on a single-trial basis (see Methods, Figure S5). We first sorted the LFP into trials that exhibited a prominent late wave (blue) and a weak late wave (gray). We extracted calcium events for these trials and used covariance analysis to quantify functional connectivity and detect cellular ensembles. **(F)** Traveling wave spatial source points in a representative mouse during stimulation (early) and late response. Note that the source points generally arise in the principal barrel (P)with a weak wave in the surround barrel (S). Note that only the principal whisker is stimulated. **(G)** Differences in wave phase-gradient magnitude across time scales of the principal barrel (ranked- sum Wilcoxon test, 764 trials across 4 mice). **(H-K)** Quantified results from functional connectivity analysis (n = 4 animals, 22 FOVs, 250 total detected ensembles). **(H)** The total number of cells within each ensemble was unchanged for ensembles associated with the late wave or a weak late wave (ns = not significant, ranked-sum Wilcoxon test). Note the small number of cells in each population. **(I)** Although the number of cells in each ensemble does not differ, we found that the number of functional connections within the ensembles is higher for weak-late wave trials, suggesting that increased sparsity supports the late wave (p < 0.001, ranked-sum Wilcoxon test). **(J)** We developed a reactivation index that quantifies how consistently cells across the ensemble are co-active. The late-wave ensembles show a significantly higher reactivation index, suggesting enhanced functional connectivity between these cells. As expected, randomly shuffling the calcium events abolishes the reactivation correlations (p < 0.001, Kruskal-Wallis with a post-hoc Dunn-Sidak test). **(K)** We quantified the weights between cellular connections in each ensemble and found that the late wave ensemble shows consistently higher weights (p < 0.001, Kolmogorov–Smirnov test).

To approach the complex task of comparing the slow, cell-specific calcium dynamics of two-photon imaging with the high-speed LFP recordings of the NeuroGrid, we developed an analysis pipeline focused on a trial-by-trial analyses of neuronal ensembles^53,54^ (**Figure 5D-E**, **Figure S5A-D**, **see Methods**). This approach overcomes the limited value of a frame-by-frame comparison of slow calcium dynamics to electrophysiology. Specifically, we sorted trials into categories based on the strength of the late wave (**Figure 5E**), which generated different wave LFP profiles – trials with a strong late wave and trials with a weak late wave. After making this distinction, we noted that the early and late response of all waves were strongly linked to the principal whisker barrel (**Figure 5F**); however, the spatial profiles of these were distinct (**Figure 5G**). Specifically, we found that weak late waves exhibited a reduced phase gradient directionality when compared to both strong late waves and early waves (**Figure 5G**, p < 0.002 ranked-sum Wilcoxon test). How are the L2/3 cellular populations altered during these changes in wave spatial dynamics? For each LFP category, we gathered all calcium events for cells in the FOV beginning 500ms before touch and ending 4 seconds after touch, used a vector-based covariance analysis to cluster cells into neuronal ensembles, and ran a trial-by-trial functional connectivity analysis on the detected ensembles (**Figure 5E**, **Figure S5A-D,** see **Methods**). These analyses allowed for the quantification of connectivity patterns in prominent ensembles that arose in trials with and without the late wave, thus enabling us to gauge L2/3 sparsity based on traveling wave spatial properties.

Across four animals and 22 FOVs at varying L2/3 depths, we detected 250 total neuronal ensembles. The average size of these ensembles was surprisingly small (<20 total cells) and did not change in trials with and without a strong late wave (**Figure 5H**, p > 0.05, ranked-sum Wilcoxon test). However, the number of functional connections in each ensemble was reduced by nearly 50% by the strength of the late wave, indicating enhanced sparsity (**Figure 5I**, p < 0.0001, ranked-sum Wilcoxon test). Despite these reduced connections, the late wave ensembles showed a dramatically increased reactivation index, a measure of the reactivity of the ensembles on a frame-by-frame basis, compared to chance (**Figure 5J**, p < 0.0001, Kruskal-Wallis test with a *post-hoc* Dunn-Sidak test). Conversely, ensembles during trials without the late wave were more stochastic, reflective of lower functional connectivity resulting in a lower reactivation index (**Figure 5J**). These results suggest an increase in the strength of functional connections in the late-wave ensembles. Indeed, neuronal ensembles associated with the late wave showed significantly increased connectivity weights, a measure of the strength of individual connections between pairs of neurons across ensembles (**Figure 5K**, p < 0.001, Kolmogorov–Smirnov test). These results directly prove that a “sparse, but strong” functional connectivity framework supports traveling waves across L2/3—partiularly during waves with heightened spatial phase gradients—and provides the first direct experimental demonstration for the sparse wave theory^7^.

### A recurrent circuit model reconciles feedback control of sparsity

How might the late wave control wS1 sparsity? Our narrowband analysis revealed nested beta-theta wave dynamics upon touch and that extended hundreds of milliseconds post-touch (**Figure 4**). Given beta is a hallmark of top-down input and goal-directed processing^50,51,55^, we turned our attention to ascertain the effect of feedback inputs on sparsity.

We hypothesized that feedback inputs could further enhance sparsity in L2/3 populations by primarily targeting inhibitory neurons. To test this hypothesis, we utilized a spiking network model, a hallmark of L2/3, operating in a state of balanced excitation and inhibition (**Figure 6A**). Synapses in the network were conductance-based. These synaptic interactions, which are considered the driving force of excitatory and inhibitory conductance, allow networks to produce robust self-sustained activity, in which the network can self-generate low-rate asynchronous-irregular activity consistent with the statistics of the spontaneous background state *in* vivo^56,57^. At scales matching the biological networks in our experimental recordings, these self-sustained states can occur with synaptic interactions in the range of 0.15 mV EPSP from rest, matching the scale of single inputs *in vivo*^57^. This means that neurons in the model experience synaptic inputs and conductance states approximating what occurs in awake, behaving animals^58–60^. The addition of local connectivity, where the likelihood of connection decreases as a Gaussian with the distance between two neurons, and distance-dependent time delays that simulate the conduction of spikes along the unmyelinated horizontal fibers connecting neurons across L2/3, results in the organization of self-sustained activity into waves that travel across the network in a systematic manner (**Figure 6B**)^2,4^. Waves in this network excite neurons as they pass over a local area in the network (gray dots, **Figure 6C**). However, we found that descending feedback projections could further increase the sparsity of these ongoing stimulus-triggered waves, creating an even more “sparse” wave, in that relatively few neurons spike as the wave propagates, a feature we observed in the experimental data (**Figure 5**). We find that predominantly inhibitory-targeted feedback occurring as an individual wave passes over a local network can substantially increase the sparsity of traveling waves in the network (**Figure 6C**). These results also hold systematically for feedback targeting different ratios of inhibitory-to-excitatory ratios (**Figure S6**). Notably, the cessation of the feedback inputs leads to a transient reduction in both the excitatory and inhibitory firing rates, reminiscent of inhibitory stabilization (**Figure 6D**)^61^. These results demonstrate a specific mechanism for how localized excitatory feedback projections onto recurrent excitatory and inhibitory neuronal networks can increase the sparsity of waves traveling across wS1 by modulating just a few neurons to spike. Such an effect could be obtained via top-down wMC feedback in wS1^25^. Indeed, silencing wMC with muscimol decreased ensemble stability in L2/3 (p < 0.0001, n = 4 mice, data not shown), suggesting wMC feedback to wS1 plays a critical role in sparsifying the L2/3 response. But the question remains, how is the prominent surface potential during the late wave produced if the population level response in L2/3 is sparsified as a function of feedback? We hypothesize that wMC feedback sculpt and modulate the late wave in wS1 by simultaneously controlling L2/3 sparsity (**Figure 5**) and gating L5 apical dendritic calcium activity (**Figure 7A**), reflected in the cortical surface potential. We examine this effect below.

**Figure 6.**
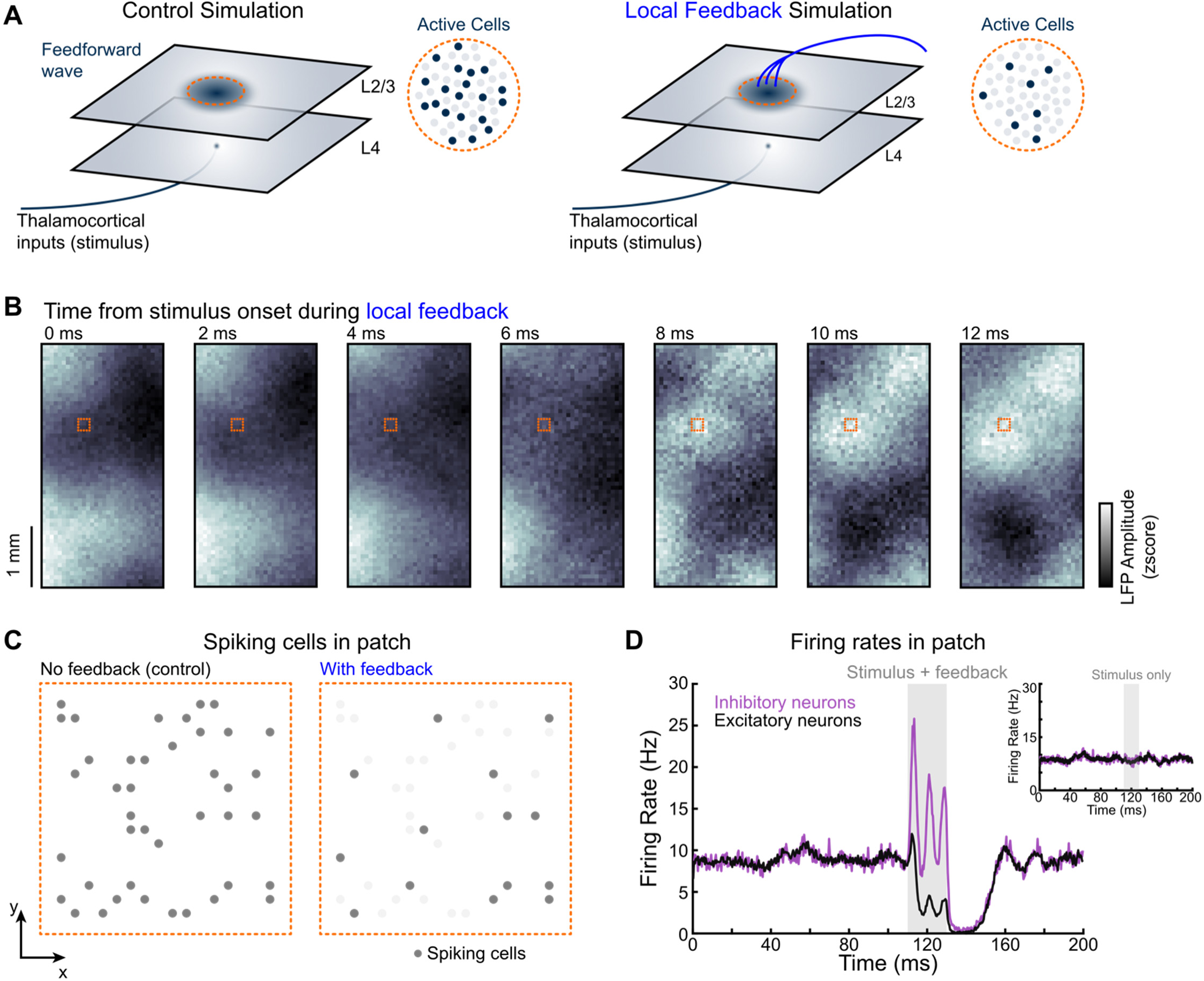
An E-I balanced model reproduces L2/3 sparsity through feedback wave-induced inhibitory stabilization. **(A)** Schematic overviewing the spiking neural network stimulation and results **(B)** Spontaneous L2/3 traveling wave LFP modeled on a 2×4 mm patch of cortex. A t = 0 ms, a sensory stimulus and feedback inputs are exerted onto a small cortical patch (dotted square box). Notice how the wave changes as a function of time. **(C)** Zoom on a 100×100 µm^2^ area of modeled cortex indicated by the dotted square box in (B) to show the responses of single cells during simulations with and without feedback **(D)** Average change in single-neuron spike rates across the cortical area with and without the feedback inputs. The feedback changes the quasi-stable fixed point in the network, which lasts for some time after the stimulation. Stimulus alone elicits a very small change in firing rate that is not readily discernible from the background (inset).

**Figure 7.**
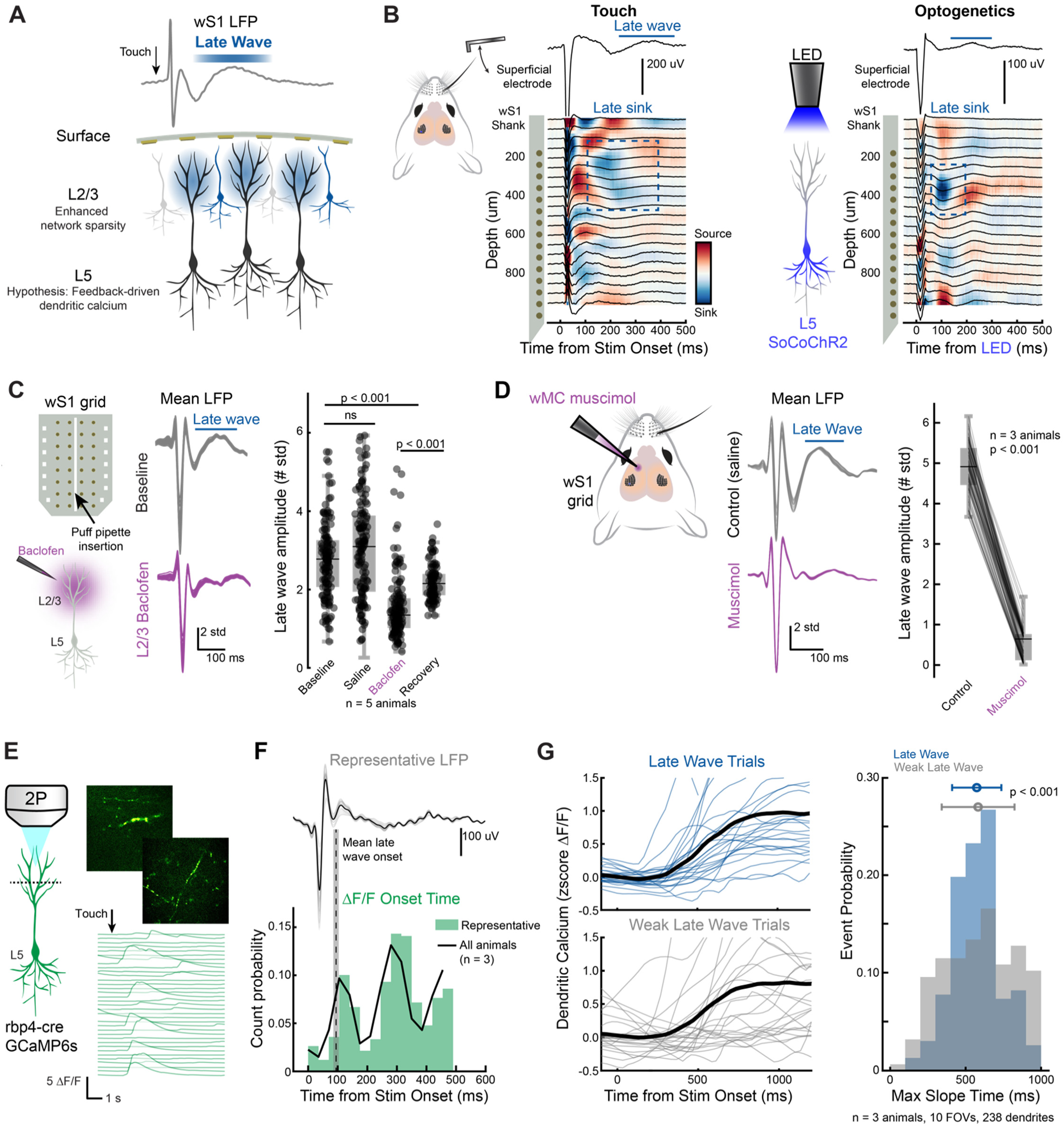
The late wave sharpens L5 dendritic calcium. **(A)** Schematic outlining the hypothesis that L5 dendrites contribute to the late wave surface potential. **(B)** Current-source density (CSD) analysis in wS1 during C1 touch and L5 optogenetic stimulation for a representative animal. (Left) Whisker touch evokes early current sinks (<50 ms) across layers, but notably, a strong delayed L2/3 sink corresponds to the late wave (∼200 ms). (Right) wS1 L5 soma-restricted ChR2 stimulation (SoCoChR2) in the same animal evokes a delayed L2/3 current sink ∼100 ms after stimulation. This sink also corresponds to a positive potential in the superficial layers (top electrophysiology trace). These results suggest an L5 contribution to the late wave, possibly originating in L5 apical dendrites. **(C)** L2/3 baclofen injections in wS1 during simultaneous surface electrophysiology. The schematic shows the surface grid design with a 50 µm wide slit through the center. We inserted a puff micropipette through the slit and injected the GABA-B agonist baclofen, which blocks dendritic calcium currents. (Center) Average surface LFP waveforms across the grid from a representative animal. Gray traces are baseline recordings for every grid channel before injections, and purple is after baclofen injection. (Right) Quantified late wave trough-to-peak amplitudes during baseline recordings, a control ACSF injection, baclofen injection, and recovery period after baclofen wash out. Each data point is a grid channel. Baclofen injections caused a significant drop in amplitude compared to the baseline and ACSF condition, and the recovery period shows a significant increase compared to baclofen. (n = 5 animals, ns = not significant, p<0.0001, Friedman test with a post-hoc Dunn-Sidak test, see **Figure S8A-C** for more results and comparisons). **(D)** wS1 NeuroGrid recording during wMC silencing with muscimol abolishes the late wave. Each line indicates a single grid channel (n = 3 animals, p < 0.001, signed-rank Wilcoxon test). **(E)** Representative touch-evoked ΔF/F transients from an apical dendrite. **(F)** Touch-evoked surface potentials and L5 apical calcium transients mapped using simultaneous electrophysiology and two-photon imaging. (Top) Representative mean surface potential during C1 touch for a representative animal. (Bottom) Dendritic calcium onset times across all animals (n = 3 animals, 238 total dendrites). **(G)** (Left) Mean touch-evoked dendritic calcium sorted into trials with and without a prominent late wave for a representative animal. Dendrites shown are the top 50th percentile of touch-responsive cells. Note the increased synchronization of dendritic calcium during the late wave trials. (Right) For all animals, we quantified the time of the max slope of touch-evoked dendritic calcium transients on a trial-by-trial basis and sorted these onset times into trials with a strong or weak late wave (n = 3 animals, 238 total dendrites, 232 dendrites showed responses during late wave trials, 157 dendrites showed responses during trials with a weak late wave). The histogram (right) shows the distribution of these max slope times along with the mean and standard deviation indicated above. While the average transient time between distributions is the same (p = 0.57), the standard deviation of the late wave distribution is significantly smaller (p < 0.001, ranked-sum Wilcoxon test following bootstrapping with 10,000 iterations to form distributions), suggesting the late wave sharpens L5 dendritic calcium transients.

### The late wave is modulated by motor feedback and L5 apical dendritic Ca^2+^ spikes

Our theoretical results (**Figure 6**) demonstrated a potential mechanism for how feedback can influence stimulus triggered traveling waves that results in the control of sparsity in L2/3 activity across wS1. If the late wave is wMC feedback modulated, our narrowband frequency analysis clearly showed an infragranular contribution to this potential via a prominent beta component (**Figure 4**). Yet, it is unlikely that NeuroGrids capture infragranular somatic activity due to the distance of the cell bodies from the cortical surface (>500 um)^62^. However, it could be possible that infragranular dendritic projections to the superficial layers—which exhibit a variety of calcium-driven regenerative spikes— contribute to the activity observed on the NeuroGrid^28^. Indeed, previous reports have suggested that somatic activity cannot fully explain delayed potentials driven by sensory input and is likely dendritic in origin^63^. In wS1, in addition to having a depolarization effect on L2/3 ^64^, studies have demonstrated the critical requirement of wMC feedback in gating L5 dendritic activity^26,47^, This is further substantiated by the fact that direct wMC inputs to wS1 synapse onto both L5 apical and somatic compartments^65^, suggesting coincidence detection, while also indirectly targeting L5 activity via the secondary thalamic nuclei (PoM)^66^. Moreover, this premise of a dendritic origin to surface potentials is also prevalent in human recordings^67^.

We began testing this hypothesis by locating the anatomical depth of the late wave with silicon depth probes and current-source density analysis (**Figure 7B**, **Figure S7A-C**). In line with previous studies^68^, we found that whisker touch drives not only an array of early sinks and sources consistent with the canonical cortical circuit, but also an extensive and delayed L2/3 current sink associated with positive potentials in the superficial layers (**Figure 7B, Figure S7A-C**). In simultaneous NeuroGrid and silicon probe recordings, we found that this delayed L2/3 sink disappeared in trials without the late surface wave, indicating the late superficial sink are tightly linked to the late wave (**Figure S7B-C**, n = 4 animals, p < 0.001, signed-rank Wilcoxon test). While the late L2/3 sink could be due to somatic spiking, we did not find a significant difference in single-unit activity firing rates in the supragranular layers across trials exhibiting a strong and weak late sink (**Figure S7D-F**, n = 4 animals, p = 0.36, signed-rank Wilcoxon test). These results suggest that superficial excitatory activity cannot fully explain the delayed L2/3 current sink. However, it is possible that L5 somatic activity drives calcium electrogenesis in superficial apical dendrites, which contributes to L2/3 current sinks^28^. Indeed, when we optogenetically activated wS1 L5 somatic activity (see **Methods**), we recapitulated a delayed supragranular current sink (**Figure 7B**). Moreover, in opposition to L2/3, L5 neurons showed an increase in single-unit firing rates during the late period in trials with the late sink (**Figure S7E-F**, n = 4 animals, p < 0.01, signed-rank Wilcoxon test). These experiments strongly implicate L5 apical dendrites in the late wave/current sink. To test their role even more directly, we pharmacologically silenced L5 calcium electrogenesis with the GABAB agonist baclofen injected 250 µm below the NeuroGrid (**Figure 7C**, **Figure S8A-C**, see Methods), which has been previously shown to primarily influence dendritic burst spiking^69^. For a period, this dramatically reduced the late wave amplitude before partially recovering after wash-out (**Figure 7C**, n = 5 animals, p < 0.001, Friedman test with a *post-hoc* Dunn-Sidak test) while largely preserving the early wave potential (**Figure S8A-C**). These experiments provide evidence that the late wave is strongly composed of signals originating from L5 apical dendritic Ca^2+^ spikes.

Consistent with this hypothesis of wMC initiated L5 dendritic spikes contributing to the late wave, we were able to abolish the late wave amplitude by pharmacologically silencing wMC with muscimol (**Figure 7D**, n = 3 animals, p < 0.001, signed-rank Wilcoxon test). Further, optogenetic inhibition of wMC with the light-gate chloride channel NpHR3 expressed in excitatory neurons (**see Methods**) also reduced the late wave amplitude (**Figure S8D-F**, n = 3 animals, p < 0.001, signed-rank Wilcoxon test). While these experiments suggest that a complex interaction between L2/3 subnetworks and L5 neurons are the origin of the wS1 late wave, they do not explain the significant delay of this large amplitude traveling potentials post touch. This variable delay (∼100-200 ms, see **Figure 2**) suggests a reverberatory circuit loop may underly this phenomenon^70^. To test this, we performed optogenetic activation of wMC with channelrhodopsin expressed in excitatory cells which drove a delayed supragranular current sink in wS1 approximately 100-200 ms after light stimulation, consistent with the delayed timing of sensory-evoked L5 apical dendritic regenerative currents (**Figure S8G**, n = 3 animals). These experiments demonstrate that the late wave is driven by wMC feedback, is tightly linked to L5 dendritic calcium spikes, and clearly visible from the surface of the brain.

### The strength of the late wave is synonymous with synchronous L5 output

How does the late wave map to L5 dendritic activity? To directly probe this relationship, we again turned to simultaneous NeuroGrid recordings with two-photon imaging during whisker touch (**Figure 7E**, **see Methods**). Dendritic calcium transients were readily measurable via transgenic lines (Rbp4- cre) labeled with GCaMP6s, and the signals were visible through the NeuroGrid (**Figure 7E**). While the decay time of GCaMP6s is slow, the transient onset time is known to be a close measure of electrophysiology onset and is sensitive to single action potentials under favorable conditions^71^. Therefore, we compared dendritic ΔF/F onset times to surface grid recordings and found the dendritic Ca^2+^ transient activation time closely tracked the late wave onset (**Figure 7F**, n = 3 animals, 238 dendrites). In fact, these data also show that L5 dendritic spike onset times can often occur even 300 ms after touch, and indeed we observe a smaller secondary positive surface potential swing on the NeuroGrid (**Figure 7F**). These results match well with recent studies in mouse visual cortex that demonstrate that visual stimuli can drive multiple low-frequency waves hundreds of milliseconds following sensory input^72^.

As with experiments in L2/3, we sorted NeuroGrid trials with simultaneous dendritic imaging into LFP that showed strong and weak late waves (n = 3 animals), where we previously found that strong late waves generated enhanced phase spatial profiles (see **Figure 5G**). In trials with prominent late waves, dendritic transients were more synchronized upon whisker touch (**Figure 7G, top**), in comparison to trials without a prominent late wave (**Figure 7G, bottom**). When we quantified the rising phase of dendritic calcium transients across animals, we discovered a significant sharpening of dendritic responses (**Figure 7G, right,** n = 3 animals, 10 FOVs, 238 dendrites, p < 0.001, ranked-sum Wilcoxon test). Our results show that wMC feedback-induced synchronization shapes traveling wave structure by sculpting a translaminar precision code. Importantly, this result also establishes how traveling waves comprise of distinct feedforward and feedback components over different timescales.

## DISCUSSION

In this work, we provide evidence that distinct translaminar circuit patterns with both cellular and subcellular origins support cortical surface traveling wave dynamics across the barrel cortex of mice. We introduced a custom, multi-modal, transparent and flexible NeuroGrid platform and dissected the translaminar circuit components underlying sensory-evoked traveling waves. Our results establish the first direct evidence that a) touch-evoked traveling waves across barrel columns comprise of distinct feedforward and feedback contributions, b) Sparse population coding in L2/3 pyramidal cells coincides with traveling wave propagation across the cortical surface, and c) traveling waves influence cortical output by shaping L5 timing. These results, in combination with our observation of wave modulation by reward-based associations, provide a link for how cortical waves correlate and carry with them information relevant to sensorimotor behaviors, stimulus representations, and sensory associations.

### Consequences of whisker-touch evoked traveling waves on coding

The local-field-potential is produced by synaptic inputs, which cause variable fluctuations in spontaneous activity and stimulus-evoked responses. These fluctuations are governed primarily by the dense recurrent connectivity of cortical networks, which induces waves traveling across cortical space^7^. Previous studies have established that these waves occur spontaneously in awake behaving conditions ^4^, and that strong driving input, such as whisker-touch, is believed to reduce variability in ongoing dynamics, producing strong correlations in spiking activity. This raises questions about the potential impact of traveling waves on evoked activity. Remarkably, our results show that strong driving inputs temporarily reset wave dynamics while still facilitating waves over a window of time, including after the stimulus has ended (**Figures 2, 3, and 4**). This result is critical as it sheds light on the possibility of stimulus feature integration based on past stimulus history and present stimulus conditions^7^. Moreover, periods with zero-phase-lag (stationary oscillation bump; no wave) could enable short-time-scale integration localized to the barrel column while periods during wave propagation could enable integration over longer time scales and across larger areas^19^, further influencing sensory integration. Additionally, based on their propagation speed, our results (**Figures 2 and 3**) suggest that these waves travel along unmyelinated horizontal fibers and have their properties shaped in a context-dependent manner (**Figure 3**). Specifically, the phase alignment across the barrel exhibited less jitter when evoked by stimuli that were associated with reward (**Figure 3F-G**). Such phase differences combined with our computational results (**Figure 6**) point to delicate changes in excitation-inhibition balance ^73^ with feedback-induced inhibition playing a key role in shaping wave-related circuit dynamics (**Figure 6 and Figure S6**)^44^. Physiologically, we demonstrate that traveling wave properties are controlled by wMC induced network sparsity in wS1 L2/3 (desynchronized asynchronous state) (**Figure 5**) and precisely timed action potentials in wS1 L5 (synchronized output) (**Figure 7**). Given that wMC feedback modulates both excitatory and inhibitory cells in wS1, and that our computational model highlights the critical role of inhibitory control on network activity^74^, suggests that traveling wave structure in wS1 reflects translaminar excitatory-inhibitory balance. Specifically, the coupling between sparse-waves in L2/3, and precisely timed dendritic regenerative spikes in L5 suggest that a unique “translaminar spacetime code” could help integrate stimuli in wS1. In particular, trials with weak late wave dynamics reflective of weak wMC-induced inhibition and increased L2/3 activity in wS1, could promote network activity dominated by fluctuations and continuous sampling. On the other hand, trials with strong late wave dynamics observed in our study, could reflect periods of optimal wMC-induced inhibition in wS1, which leads to higher reliability and a reduction in noise correlations ^75^ (**Figure 3, 5, and 7**)

### Implications of a translaminar L2/3 to L5 spacetime code

Barrel cortex neurons are known to exhibit selectivity to the global direction of the tactile stimulus^76^. This would imply that individual neurons integrate across space to extract higher-order stimulus features. It would thus seem apparent that TWs elicited via touch from one whisker could prime surrounding cortical columns to upcoming stimuli by signaling their reference frame to adjacent somatotopic spaces in the cortex, enabling more collective integration of stimulus features rather than local independent variables^46,77^ . Critical to this facet are neurons in the supragranular layers of the cortex that play a key role in orchestrating wave propagation (**Figure 5, and Figure 6**). Here, excitatory layer 2/3 (L2/3) neurons are known to exhibit mixed selectivity with a sparse representation of touch and object location^78^. These sparse L2/3 patterns likely require feedforward and recurrent intracortical connections^11^. Through these intracortical circuits, L2/3 ensembles may integrate tactile features that span multiple whiskers ^46^. We hypothesize that TWs enhance the fidelity of sensory-evoked L2/3 ensembles via sparse network activation^79^. These ensembles, possibly gated^73^ dynamically by traveling waves, could facilitate the representation of certain stimulus features, improve dynamic range, and aid the classification of multi-whisker stimuli. The transformation of the feed-forward stimulus in L2/3 should enhance the robustness of feature classification to noise and variations of the input signal to facilitate downstream processing^80^. In addition to direct feed-forward processing, traveling waves controlled via local and long-range recurrence, could help link stimulus features across time and contextualize stimuli based on past information.

The effect of traveling waves on translaminar circuits beyond L2/3 is an open question. Studies in brain slices suggests that layer 5 neurons play a role in controlling superficial excitation^81^ and the spread of traveling waves^82^. The microcircuit that enables this translaminar dynamic likely plays a critical role in shaping cortical output and behavior. Moreover, L5 neurons not only receive direct feed- forward drive from L2/3 but also top-down motor feedback onto their apical dendrites. It is thus feasible that traveling waves generated via bottom-up thalamic drive, could be organized by translaminar interactions sculpted by long-range feedback. Our recordings support this theory and revealed in addition to the sensory-evoked early wave, a late wave lasting hundreds of milliseconds post stimulus. The strength of the late wave was time-locked to L5 dendritic Ca^2+^ spikes, nearly abolished by GABAB agonists injected into the calcium rich zone, and strongly modulated by wMC, signifying a role for feedback-driven wave control (**Figure 7**). Moreover, L2/3 neurons alone did not exhibit strong modulation during periods of the late wave (**Figure 5 and Figure S7E**). This finding asserts that the recorded late wave surface potentials are a reflection of regenerative dendritic spikes occurring a few hundred microns below the pia^83^ – events which are tightly linked to sensory perception^27^, and consciousness^84^. Notably, such large surface potential swings are in line with previous reports^28^, and is supported by the fact that the dipoles created by the distinct geometry of pyramidal cells have long been considered to dominate field potential recordings captured using EEG and MEG^85,86^

Overall, our results suggest that traveling wave dynamics from the cortical surface reflect distinct translaminar circuit features critical for optimal sensory perception.

### Motor feedback is critical for organizing traveling waves via changes in E-I balance

Sensory perception is an active process where the brain receives and processes information based on previous experiences, expectations, and current goals. Here circuits bind and route information across specific pathways over behavioral timescales to create perceptual experience.

Within this context, the interaction between sensory and motor cortices is crucial in accurate sensory perception and discrimination and controlling what sensory information the brain will receive^87^. In the barrel cortex, interactions between sensory and motor cortices are of fundamental importance for accurate sensory perception and discrimination^47,48,88–90^. wMC projections show a strong preference for deep layers (L5/6) and L1, but are known to also broadly engage different excitatory and inhibitory cell types in sensory cortex^25,65^. Pure excitatory motor cortex feedback to wS1^90^, has been reported to enhance spiking responses to touch via supralinear amplification^91^, encode movement and touch responses^88,89^, drive spiking across cortical layers^47^, and generate calcium transients in L5 apical dendrites^26^. Inhibitory motor feedback effects on the contrary has been reported to suppress responses to distractor stimuli in barrel cortex^48^.

These sensory-motor interactions occur over timescales of 10’s to 100’s of milliseconds, suggesting that connections are highly recurrent and not purely monosynaptic. More importantly, it suggests that traveling waves might be able to bind several antecedents, along with past history, and current stimulus conditions. Our data supports a model in which spatially restricted motor feedback to wS1 targeting both excitatory and inhibitory neurons enhance sparse coding in L2/3; “sparsifying” the wave (**Figure 6**). This is confirmed by our computational results which show mechanistically how this phenomenon can emerge across a wide range of feedback-targeted inhibitory and excitatory ratios (**Figure S6**), consistent with recent results from recurrent circuit models signifying a change in the stability point of the network^11^. Notably, we hypothesize that such feedback induced changes in wave patterns might be a way to selectively engage ensembles, a form of selectivity within mixed representations^52^.

Given wMC to wS1 pathway terminates on layer 5 and layer 2/3 neurons (Mao et al., 2011), alludes to a critical role of wMC feedback, not only in sparsifying L2/3 but also sharpening L5. This result is particularly significant as wMC feedback has been shown to elicit calcium transients in L5 apical dendrites, a feature critical for accurate sensory perception^27^. Our data add to this framework, demonstrating that not only can dendritic potentials be organized into a traveling wave, but also show that wMC feedback sharpens L5 dendritic calcium transients. Such an effect can induce synchronized burst-firing in L5 somas, leading to heightened downstream transmission for goal-directed sensory discrimination. Our results are also in line with the retinotopic mapping of cortico-cortical feedback circuits recently discovered to engage apical dendrites in visual cortex^72^, suggesting a generalized role of reverberating waves and dendritic excitability.

Our results are also in line with recent works supporting a translaminar code^92,93^ in which inhibitory microcircuits mediate L2/3-L5 coupling. Specifically, somatostatin (SST) interneurons were shown to be responsible for L5 suppression of touch-evoked activity during optogenetic activation of L2/3^92^, highlighting gain control. Other reports have shown that that deep-layer SST interneurons are potent modulators of L5 apical dendritic activity^93,94^. These findings mirror our observations of sparse L2/3 maps sharpening L5 during passive touch in a traveling wave format, which leads to speculate that traveling waves across superficial layers might depend on unique interneuron subtypes.

### Outlook

Our study shows that touch-evoked traveling waves are supported by a unique translaminar spacetime circuit pattern. We utilized NeuroGrids, two-photon imaging, silicon probes, pharmacology, optogenetics, and network modeling to establish a connection between the dynamics of traveling waves and the underlying circuit dynamics. Using our custom-fabricated grids, we first showed that whisker touch evoked prominent patterns of traveling waves across the mouse barrel cortex (**Figure 1,2**). Our data suggest these touch-evoked broadband waves can be classified into two primary categories: early-onset feedforward waves in the beta-gamma frequency bands and late-onset, feedback-driven waves in the beta-theta bands that are modulated by behavioral context (**Figures 2, 4, 7**). In agreement with previous computational efforts describing how traveling waves shape the cortical landscape, we found that different traveling wave patterns are hallmarked by changes in L2/3 network sparsity (**Figure 5**), establishing the first experimental demonstration of the “sparse wave” theory. We computationally validated these results using a recurrent balanced state spiking neural network model, which revealed that feedback inputs to even a small subpopulation of inhibitory cells within the network could have widespread effects in cortical sparsity (**Figure 6**), reflecting a state of inhibitory stabilization. These results led us to question how a prominent late wave surface potential emerges as sparsity in L2/3 constricts superficial activity. We show that the late wave dynamics largely originate from calcium electrogenesis in L5 apical dendrites (**Figure 7**), also establishing a biomarker of dendritic dynamics that one can reliably map from the brain surface. In summary, we found that whisker touch evokes an early-onset feedforward wave, a motor-feedback-driven late wave, and that this feedback initiates inhibitory action to concomitantly shape L2/3 population sparsity and drives precise L5 dendritic potentials, thus inducing a translaminar spacetime code.

### Limitations of the study

In this work, we did not distinguish between L5 subtypes in contributing to wave dynamics. It is conceivable that wMC feedback sharpens and synchronizes activity between sublayers, heightening both cortico-cortical and subcortical signaling. Another strong possibility is that wMC feedback synchronizes calcium spikes across L5B cells to promote descending activity, which has previously been shown to be a critical component for sensory perception^27^ . In fact, thick-tufted L5B pyramidal neurons show enhanced bursting during active touch in addition to a secondary volley of spiking 50- 120 ms following touch^95^, similar to the timescales of the late wave we observed. Critically, one could envision a cortico-thalamocortical loop in which traveling waves gate timing in L5B, which feeds to the thalamus, and in turn feeds back to L5A. Given anatomical evidence that L5A synapses onto L5B dendrites^18^ as well as L2/3^96^, could set the stage for unraveling precise phase-locking across thalamocortical loops, and elucidating the role of L5 in controlling superficial wave spread^82^.

## ACKNOWLEDGEMENTS

This work was supported by the following grants to K.J. NIH R21EB029740 Trailblazer Award; Human Frontiers Science Program (HFSP) RGY0069; ORAU Ralph E. Powe Junior Faculty Enhancement Award, Purdue Institute for Integrative Neuroscience, and the National Institutes of Health, New Innovator Award NIH DP2MH136494. This material is also based upon work supported by the Air Force Office of Scientific Research under award numbers FA9550-22-1-0078 and FA9550-23-1-0701. Any opinions, findings, and conclusion or recommendations expressed in this material are those of the author(s) and do not necessarily reflect the views of the United States Air Force. D.L.G. was supported by the HHMI Hanna Gray Fellowship and the Burroughs Wellcome Fund Postdoctoral Enrichment Program. H.F.K. is supported by the NIH T32 – 5T32DC016853 and NSF GRFP (DGE-1842166).

## AUTHOR CONTRIBUTIONS

D.L.G fabricated grids, analyzed data, designed, and performed experiments. H.F.K performed detailed calcium analysis, wave analysis, and assisted with result interpretation. H.V.S.K performed the active touch behavior and experiments. S.Y designed and built the microscope. C.S and L.E.M designed and carried out large-scale network simulations along with interpretation. S.R.P and K.J designed experiments. S.R.P and K.J provided guidance. K.J conceived and led the overall project and acquired funding to carry out the study. D.L.G and K.J wrote the paper with inputs from all authors.

## RESOURCE AVAILABILITY

### Lead Contact

Requests for further information should be directed to and will be fulfilled by the lead contact, Krishna Jayant.

### Materials Availability

This study did not generate any new reagents. We do not have plans to broadly disseminate NeuroGrid surface arrays but interested readers can direct their inquiries to the lead contact.

### Data and Code Availability

Data and code generated in this study will be uploaded to a public repository upon manuscript acceptance.

## EXPERIMENTAL MODEL AND SUBJECT DETAILS

### Animal subject details

All experimental procedures were conducted in accordance with the guidelines set forth by the NIH, Purdue Institutional Animal Care and Use Committee (IACUC), and the Purdue Laboratory Animal Program (LAP). All experiments were conducted in adult mice ages 3-8 months old. For passive touch recordings, we used mice with a C57BL/6J (The Jackson Laboratory, #000664) background kept on a 12-hour light/dark cycle in conventional housing with unrestricted access to food and water. Male and female mice were used in approximately equal numbers. For L2/3 imaging experiments, we used Thy1- GCaMP6s transgenic mice (Jackson, C57BL/6J-Tg (Thy1-GCaMP6s) GP4.3Dkim/J, #024275). For experiments with targeted L5 expression, we used Rbp4-Cre mice (MMRC, B6.FVB(Cg)-Tg(Rbp4- cre)KL100Gsat/Mmucd, #037128-UCD). For active-touch experiments, we used EMX1 mice (Jackson, B6.129S2-Emx1tm1(cre)Krj/J mice, #005628) outcrossed with CD-1 mice (Jackson, #022) for multiple generations. We used both sexes maintained in reverse light-dark cycle (12:12 hr.). Behavioral training and recordings were conducted during the animal’s subjective night.

## METHOD DETAILS

### Grid microfabrication

We used conventional photolithography approaches to fabricate 10-15 parylene-based surface grids simultaneously on 4” silicon (Si) wafers. We first cleaned the Si wafers (Nova Electronic Materials) by sonicating in toluene, acetone, and IPA for 5 min each. We then dried the wafers with nitrogen and performed a 1 min O2 plasma clean (March Jupiter II). For parylene deposition, we used chemical vapor deposition (Specialty Coating Systems) and 3 g of dimer to deposit ∼3 µm of parylene C onto 4-5 wafers simultaneously. To remove leftover monomers from the parylene film, we vacuum baked the wafers overnight at 140 ^°^C.

To form the metal recording sites, leads, and connecting pads, we used photolithography. We spun 300 nm of LOR 3A and baked for 10 min at 150 ^°^C followed by and 3 µm of AZ1518 and a 2 min bake at 100 ^°^C. We exposed the photoresist with the first metal pattern with a power of 224 mJ/cm^2^ using a maskless aligner (Heidelberg MLA 150) and developed the wafers for 2.25 min in MF-26A followed by a water rinse. Following development, we performed a brief 10 s O2 plasma clean (March Jupiter II). For metal deposition we used RF sputtering (PVD SD-400) to deposit 300 nm of Au with a 5 nm Ti adhesion layer. We used deposition rates of ∼1.4 nm/min for Ti, 5 nm/min for the first 20 nm of Au (this reduces thin-film stress), and 7 nm/min for the rest of the Au layer. We performed liftoff overnight in Remover PG and rinsed with acetone and IPA. To drive off any leftover solvents, we baked the wafers at 110 ^°^C for 5 min.

To reduce the electrochemical impedance of the recording electrodes, we repeated the above photolithography steps with a different mask pattern to deposit nanoporous Au onto only the recording sites^31,32,97^. For this sputtering process, we co-sputtered Au and Ag simultaneously at DC powers of 50W and 120W, respectively, to a final metal thickness of 400 nm. Following liftoff, we de-alloyed the metal film in 70% nitric acid for 3 min at 70 ^°^C. This process etches the Ag in the alloy and leaves behind porous Au. We rinsed the samples in water, dried them with nitrogen, and dehydrated them at 70 ^°^C for 3 min and 110 ^°^C for 5 min.

To encapsulate the deposited metal, we performed a brief 30 s O2 plasma clean to activate the surface of the bottom parylene layer, deposited another 3 µm film of parylene C with the same technique as above, and vacuum baked overnight at 140 ^°^C for a final parylene thickness of 6 µm. We found this ultra-thin film to be difficult to handle post-fabrication. Therefore, we performed another photolithography step to strengthen the backend of the probe with a layer of 15 µm SU-8. To do this, we again activated the parylene surface with a 30 s O2 plasma clean, spun SU-8 2010 onto the surface to a thickness of 15 µm, and baked the wafers for 3 min at 95 ^°^C. We used a mask aligner (MA6) to expose the resist with a dose of 160 mJ/cm^2^, baked the sample for 3 min at 95 ^°^C, developed in SU-8 developer for 3 min, and rinsed with IPA. To reduce SU-8 stress and improve the SU-8/parylene adhesion, we hard baked the wafers with a ramping bake (2 min at 100, 110, 120 ^°^C, and 1-2 hr at 130^°^C). Importantly, this SU-8 layer only supports the backend of the probe (i.e., the large connecting pads and approximately 75% of the probe length) while the grid array portion that sits on the brain remains only parylene and ultra-flexible. At no point have we observed delamination between the parylene-parylene or parylene-SU-8 layers, which we attribute to the O2 surface activation steps, vacuum baking steps, and temperature ramping.

The final fabrication step on the Si wafer is to etch away parylene to shape the probe and expose the porous Au recording sites and large Au connecting pads^98^. A 3 µm parylene etch was necessary to expose the recording sites and a 6 um parylene etch needed to define the outer edge of the probe. We found the simplest approach was to perform the etch steps simultaneously. We spun AZ9620 to a thickness of 15 µm, baked for 4 min at 100 ^°^C, exposed with maskless lithography (2100 mJ/cm^2^ exposure), and developed for 15 min in MF-26A. To avoid over-etching the recording sites, the mask holes above the recording electrodes were 10 µm in diameter, which expanded to a final diameter of ∼25 um during the etch process. We used a high-power (250 W) O2 plasma supplemented with Ar and SF6 to etch the parylene layers (March Jupiter II). Importantly, we avoided overheating of the sample by etching for 1 min followed by 1 min cool-down period of no plasma. We repeated this cycle 6-8 times for a total etch time of 6-8 min until the Si substrate was visibly clear of parylene. Finally, we stripped the etch mask in acetone and rinsed the final wafer product in acetone.

### Grid packaging

To remove the parylene probes from the Si substrate, we submerged the entire wafer in DI water and used fine-tip surgical tweezers to gently separate the parylene from the underlying wafer. To facilitate this process, the backend of the probe has a “handle” with only parylene and SU-8 layers. This handle allowed us to begin the separation of the parylene and silicon without damaging important metal structures. In our experience, once the separation begins, the parylene probes cleanly separate from the Si substrate. We also note that the SU-8 thickening layer facilitates parylene removal and found it difficult to remove probes constructed of only 6 µm parylene. Furthermore, we anecdotally note that probe removal soon after fabrication (< 24 hr) dramatically reduced probe curling.

After removal and drying, we bonded the backend of the probes to a laser-cut 0.010”-thick polyether-ether-ketone film^30^ (CS Hyde, 37-10F-24“x24”) using a 20% mixture of poly-vinyl alcohol in water (Sigma, 341584-25G). This film allowed for stable insertion of the probe backend into a 32 Ch ZIF clip (Hirose, FH12-32S-0.5SH (55)), surface mounted to a custom 1.0×0.5” printed circuit board. Onto the PCB we also mounted a ground pin and 32 Ch Omnetics connector (NPD-36-VV-GS), which allowed us to interface the parylene grids with a 32 Ch Intan amplifier (RHD 2132).

### Surgical procedures

Mice were first deeply anesthetized with 3 to 4% isoflurane. During surgery, anesthesia was maintained at 1 to 1.5% isoflurane with an oxygen flow rate ∼0.1 L/minute. An infrared warming pad (Kent Scientific) was used to maintain the body temperature. Carprofen and Dexamethasone (0.6mg/kg body weight) was injected subcutaneously, and lidocaine injected under the scalp after the induction of anesthesia. Eye ointment was applied, and the scalp shaved and sanitized before the incision. After scalp removal, we briefly applied 3% hydroperoxide to remove excess tissue and immediately cleaned the skull with saline perfusion. Vetbond (3M) was used to seal the tissue surrounding the skull region. We then used dental cement to adhere a custom-designed titanium headplate (Parkell) to the skull centered over the left barrel cortex (1.5 mm posterior to bregma and 3.5 mm lateral to midline). Once the cement solidified, we used a dental drill to partially thin an ∼5 mm area of bone around the barrel cortex until the vasculature was clearly visible when soaked in saline. We then dried the skull, sealed the exposed bone with Vetbond, and covered the entire area with Kwik Cast silicone (World Precision Instruments).

Animals were given 3-7 days to habituate to the headplate. 1 day prior to recordings, we trimmed all but the C1 whisker and performed intrinsic optical imaging to locate the C1 barrel (described below). On the day of recordings, the anesthesia and subcutaneous injections described above were performed again. For a ground electrode, we performed a small craniotomy in the forebrain and cemented in place an Ag/AgCl wire inserted between the dura and skull. To access the barrel cortex, we used a dental drill to perform a 2-3 mm cranial window (surrounded by and additional ∼2 mm region of thinned skull) centered on the C1 barrel. We regularly perfused the surgical area with clean ACSF. For all grid recordings, we also performed a durotomy with a dura hook (Fine Science Instruments, 10032-13). For experiments requiring only a silicon probe, the cranial window was <1 mm in diameter with a 2-3 mm area of thinned skull around the window. Upon completion of the surgery, we covered the cranial window with a small piece of surgical foam (Surgifoam) soaked in ACSF and transferred the animal to the recording rig where it remained head-fixed for the duration of the experiment.

### Grid recordings and electrophysiology

To place the grids over the C1 barrel, we mounted the fully-packaged probe (grids, custom PCB, head- stage) onto a manual micromanipulator. After the craniotomy and durotomy, we transferred the animal to the recording rig for head-fixation. While still lightly anesthetized, we removed the surgical foam and filled the area above the cranial window with ACSF. We then placed the grids on the ACSF, which were held in place through surface tension. Slowly, we aspirated the ACSF, while adjusting the micromanipulator to maintain the grid position over the barrel. Upon initial contact we with brain, we found that the grids did not strongly adhere and could easily be removed through the addition of more ACSF. To promote adhesion, we maintained a very thin film of ACSF over the grids and brain, aspirated the majority of the ACSF (without completely drying the brain), re-wetted the surface with another thin film of fresh ASCF. We repeated this process every ∼5 min over the course of ∼30 min. We found that this simple protocol led to strong adhesion between the grids and brain such that a pool of ACSF no longer lifted the grids off the brain. For experiments where only surface electrophysiology was required, we placed surgical foam soaked in ACSF over the grids. For experiments requiring grids in combination with pipettes or silicon probes, we maintained a layer of ACSF over the cranial window for the duration of the experiment and gently replenished this solution every ∼30 min. Finally, for experiments requiring grids with two-photon imaging, after placing the grids onto the brain surface, we aspirated ACSF to a thin layer, pressed a 3 mm coverslip (Warner Instruments, 64-0720) onto the grids and brain with a micromanipulator, and cemented in place. In most cases, we carefully cemented only the areas of the coverslip not in contact with the grids (see **Figure 1** schematic), which allowed us to recover the parylene probe after recordings.

All electrophysiological data was acquired with Intan Technologies head-stages (32 and 64 Ch) interfaced to an RHD Recording System using RHX Data Acquisition Software. The sampling rate was 20 KHz. For grid recordings with simultaneous depth recordings, we used Cambridge Neurotechnologies H5 64 Ch linear probes (800 um electrode span, 25 µm vertical spacing between electrodes) or H8 64 Ch two-shank probes (250 µm shank spacing, 930 µm electrode span, 30 µm vertical spacing between electrodes). These ultra-thin silicon probes were optimal for insertion through the 50 um-wide slit through the center of our surface grids. For experiments requiring only silicon probes recordings, we used UCLA 64D silicon probes electroplated to an electrochemical impedance of ∼300 KΩ (1.050 mm electrode span, 25 µm vertical spacing between electrodes).

### Viral injections

We performed injections 4-6 weeks in advance. All anesthesia and surgical protocols were identical to the above protocols. However, we made a small incision in the scalp rather than completely removing the skin. We then made a small burr hole in the skull using a dental drill. An injection micropipette (Sutter BF100-50-10, Sutter P-1000 puller) with a diameter of <50 µm and backfilled with mineral oil was mounted to a microinjector (World Precision Instruments, Micro4 controller, UMP3 micro-syringe pump) for injections. For all injections, we inserted the pipette ∼100 µm past the target depth, then slowly raised the pipette up to the target depth. All injections were performed at a rate of 20 nL/min and we waited 5-10 min after injection completion before slowly removing the micropipette. For wS1, we used stereotactic coordinates 1.5 mm posterior to bregma and 3.5 mm lateral to midline. For wMC, we used stereotactic coordinates 1 mm anterior to bregma and 1 mm lateral to midline. Following injections, we sutured the scalp and allowed the animal to recover in a heated cage. The AAV, titer, volume injection, and injection depth are in **Table 1**.

**Table 1.** Virus injection details.

| Experiment | Animal line | Plasmid | Addgene# | Titer (vg/ml) | Volume (nL) | Depth (μm) |
| --- | --- | --- | --- | --- | --- | --- |
| wS1 L5 apical dendrite imaging | Rbp4-cre | pAAV-hSyn1-Flex-mRuby2-GSG-P2A-GCaMP6s-WPRE-pA | 68720 | $1 \times 10^{13}$ | 250/depth | 600, 500 |
| wS1 L5 ChR2 stimulation | Rbp4-cre | pAAV-hSynapsin-FLEX-soCoChR-GFP | 107712 | $1.0 \times 10^{13}$ | 500 | 500 |
| wMC ChR2 stimulation | C57BL/6 | pAAV-CaMKIIa-hChR2(H134R)-mCherry | 26975 | $2.30 \times 10^{13}$ | 100/depth | 800, 600, 400, 200 |
| wMC NpHR inhibition | C57BL/6 | pAAV-CaMKIIa-eNpHR 3.0-EYFP | 26971 | $1.0 \times 10^{13}$ | 350/depth | 600, 300 |

### Intrinsic optical imaging

1-2 days before recordings, mice were anesthetized with 3 to 4% isoflurane, and placed under a CMOS camera (Blackfly USB3, Teledyne FLIR) equipped with a DLSR macrolens (Nikon). If necessary, we further thinned the skull region. The camera and lens were angled for a perpendicular visualization of the barrel cortex. We maintained a light anesthesia for the animal (∼0.75% isoflurane) and a 2X higher oxygen flow rate than our cranial window surgical procedures (∼0.2 L/min) to maintain relatively high blood oxygenation. An infrared heating pad was used to maintain the body temperature during anesthesia. We filled the skull area with mineral oil and used a coverslip to planarize the surface for a stable image intensity. The thinned, soaked bone was transparent to the point of easily visualizing the underlying vasculature. We used a multi-wavelength LED for all imaging (Lumencor, Spectral X). We trimmed all whiskers other than the C row and placed a small metal hook over the C1 whisker. The hook was attached to a stepper motor (NEMA 17 motor, DM556 motor driver) controlled by an Arduino UNO for rapid vibration at 30 Hz. An image of the vasculature was taken as a reference with green LED illumination. A NIR LED was used to illuminate the thinned scalp for intrinsic imaging. Custom-written MATLAB (MathWorks) and Arduino codes were used for controlling the experiment. During each imaging trial, we recorded 100 frames of baseline data (25 fps) followed by an additional 100 frames during whisker stimulation. The frames taken before and during whisker deflection were averaged separately and the difference between the two averaged images formed the intrinsic optical signal for one trial. The trial was repeated for 6 to 15 times, and the resultant image from each trial was averaged until the contrast of the image was clear and the barrel distinguishable from the background. Using ImageJ, we smoothed the final image with a gaussian filter and overlaid the intrinsic signal with the vasculature reference.

### Passive whisker stimulation

Awake, head-fixed animals were allowed to run on a wheel, but were largely stationary due to the lack of incentive for locomotion (**Figure S2F**). For recording touch-evoked potentials, we performed passive stimulation of the C1 whisker with a metal pole driven by a motor that fully brushed past the whisker rapidly (NEMA 17 motor, DM556 motor driver) once during a trial. Whiskers were not contacted again and mice freely whisked after stimulus presentation. We used a trial-to-trial interval of 10-20 s. All experimental systems were synchronized with an Arduino microcontroller (Arduino UNO).

### Active-touch: Training and recordings

For the active-touch paradigm, we replicated previous work demonstrating a pneumatic system that extends pistons into the whisking range of head-fixed animals^46^. The organization of a single trial can be found in (**Figure S3A**). Briefly, animals must reach a running threshold of 5.6 cm/s before the Go or No-Go stimulus extends into the whisking range for an active sampling period of 1.3 s. For the Go stimulus, the animal must lick within this window. If successful, a water reward is delivered and an inter-trial interval of at least 4 seconds is used. For the No-Go stimulus, animals must withhold a lick. If the animal correctly rejects the No-Go, an inter-trial interval of 4 s is used. If the animal licks, the trial is considered a false alarm and an inter-trial timeout of 14 s initiates. To control the behavioral state of the animal, before the next trial initiates, the animal must again reach the running threshold. Throughout training and recordings, we used white noise to drown out sounds from the pneumatic parts and water delivery system.

For discrimination training, we began with the headplate surgery. Animals were given 2 to 3 days for headplate habituation and recovery. Mice were then head-fixed and allowed to habituate to the running wheel for 3 to 4 days for one hour every day. These mice were then water restricted and their whiskers were trimmed to leave C1 and D1 whiskers on the right whisker pad. We then introduced the Go stimulus (C1 piston) and trained animals on classical conditioning for 3 to 4 days to form the association between C1 touch and water reward. After this period, we introduced the No-Go stimulus (D1 piston) until mice showed anticipatory licking for the Go stimulus (3-4 days). Finally, mice were moved to operant training where they learned to withhold their licking for the No-Go stimulus (>5 days). Training every day for 1 hour was maintained until these mice reached a discrimination index, *D*′, greater than 1.5 for at least two consecutive sessions (**Figure S3B**). We calculated *D*′ with the following equation

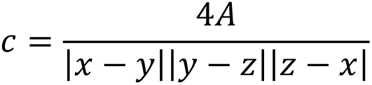

Where Z is the inverse cumulative distribution function of gaussian distribution.

1-2 days before recordings, the C1 and D1 whisker barrels were identified through intrinsic imaging. On the day of the electrophysiology recording, mice were briefly (20 - 30 minutes) anesthetized to perform a craniotomy over the identified barrels. The exposed area was covered with silicone gel (Dowsil) and Kwik Cast silicone (World Precision Instruments). We allowed animals a minimum of 3 hours of recovery before beginning recordings. For electrophysiology during the discrimination task, mice were placed in the experimental rig and grids were lowered over the Go and No-Go barrels to record neural activity while the mouse discriminated between the presented stimuli. We performed recordings in 5 animals. However, one animal was removed from the discrimination analysis due to an experimental issue that led to falsely detected licks. The data for this animal was included when analyzing the effects of whisking and touch on traveling waves (**Figure S3C-D**), but was removed for analyses comparing Hit and Correct Rejection trials (**Figure 3, Figure S3F**).

### Active-touch: Whisker tracking & kinematics

A high-speed infrared camera (Photon focus DR1) was used to track whisker kinematics at 500fps during the recording session. Camera video frames were recorded on Intan 512 controller and synchronized with neural data via external triggers generated by a National Instruments card. For post processing, DeepLabCut (DLC) was used to track whisker movement and obtain touch times^49^. For the DLC training data set, we manually labelled 200 video frames for each animal. Four labels were evenly spaced to mark each whisker in a video frame. The DLC neural network was trained for 360k iterations, and the final labels were manually checked for accuracy. The whisker position was calculated for each label for both the whiskers with reference to a user defined point on the whisker pad. Whisker phase was bandpass filtered between 1-30Hz. The whisker bend was calculated from the three distal labels on each whisker using Menger curvature (*c*).

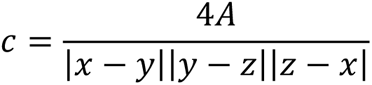

Where *x*, *y*, *z* are the coordinates of three labels on a whisker, and *A* is the area of the triangle formed by these three labels. Whisker touch frames were identified from when the whisker label enters a region of interest (manually marked around the touch surface).

### Active-touch: Lick detection

During training the headplate of the mouse was connected to 5V of the Arduino board, and an analog input pin (with a pull-down resistor to ground) connected to the water spout was used to read changes in voltage when the mouse licked. During electrophysiology, to avoid lick induced artifacts in the recording quality, the analog signal from a piezoelectric bender (BA3502, Micromechatronics, Inc.) attached to the lick port was used as lick signal. This signal was amplified by an operational amplifier and was sent in real time to an Arduino microcontroller and Intan 512 controller. The signal was synchronized with neural data via external triggers generated by the National Instruments card. Lick signal was considered a lick if it crossed a threshold of 2 volts.

### Two-photon imaging

Two-photon imaging was performed using a laser-scanning microscope (Sutter MOM, Sutter Instruments) fitted with resonant scanning mirrors (Vidrio Tech). We used a tunable femtosecond pulsed laser (Insight X3, Spectra Physics) for the excitation source with beam intensity controlled with an electro-optical modulator (Conoptics, model 350-50). We used a Nikon 16X/0.8NA water-immersion objective with a 3 mm working distance and imaged at wavelengths of 940 nm with powers of <55 mW at the focal plane. All experiments were controlled with ScanImage software with a 512×512 pixel field of view sampled at 30 Hz. To locate the grid on the brain surface without damaging the metal or parylene, we used low powers (<5 mW). We set the z = 0 focal plane to the grid center, then lowered the objective to the depth of interest before increasing the laser power. For L2/3 imaging, we performed touch trials at multiple depths between 150-275 um at 1.5X magnification with 30-50 trials per depth and a minimum spacing between focal planes of 25 µm. For dendritic imaging, we used a 3-5X magnification and imaged at depths between 50 µm (i.e., tufts) and 200 µm (apicals).

### Optogenetics

All optogenetic light stimulation was performed with a fiber-coupled LED (200 um diameter core) placed on either the surface of the brain or surface of thinned skull. ChR2 was activated with a 470 nm LED (Thorlabs, M470F3, 7 mW) and NpHR with a 595 nm LED (Thorlabs, M59F2, 4 mW).

For wS1 L5 ChR2 activation (**Figure 7B**), the LED was placed at an approximately 35° angle to the silicon probe. We reduced the light artifact on the silicon probe by facing the recording electrodes away from the light direction and offsetting LED stimulation by ∼200 um laterally from the probe insertion sight so that the maximum light intensity did not hit the backside silicon. For these experiments we also used the analog output of an Arduino to ramp the LED power to reduce the photo artifact^99^. We ramped the LED power up for 15 ms to the maximum power, held the maximum power for 5 ms, then ramped down for an additional 15 ms for a total stimulation time of 35 ms. This method reduced, but did not abolish the light artifact. We used a trial-to-trial interval of 10 s.

For wMC ChR2 activation during wS1 silicon probe recordings (**Figure S8G**), the LED was placed perpendicular to the brain surface and brief square-wave pulses of 1, 5 or 10 ms delivered. We used a trial-to-trial interval of 10 s. For wMC NpHR inhibition (**Figure S8D-F**), the LED was placed at an approximately 35° angle to the surface of wMC, which allowed us to vertically insert a silicon probe in a subset of animals to validate single-unit inhibition. For these experiments, we turned on the LED 1 s prior to whisker touch and the light remained on for another second after touch. We interleaved trials with the LED on and off to observe the effects of wMC inhibition on the late wave. We used a trial-to-trial interval of 10 s.

### In vivo pharmacology

For in vivo pharmacology, we pulled glass micropipettes to a diameter of ∼20 um and attached the puff pipettes to a pneumatic microinjector (Applied Scientific Instrumentation, MPPI-3). All injections were performed with a series of 10 ms puff pulses spaced 1-2 seconds apart. For wS1 baclofen (**Figure 7C**), we placed grids over the C1 barrel and navigated the puff pipette to the small opening in the grid center. We used a 50 µM baclofen solution dissolved in fresh ACSF and injected 20-50 nL at 10 nL/min at a depth of 250 µm. Immediately following injections, we recorded data for 30 min with a trial-to-trial interval of 15 s to observe the effects of baclofen. Data recorded 30-90 min post injection was considered the “recovery” period, when we observe the effects of baclofen to dissipate. We conducted baseline recordings prior to baclofen injection. As a control, we injected ACSF at the same depth and rate either before the baclofen injection or after the recovery period. For wMC muscimol (**Figure 7D**), we used a 5 µg/uL solution (Sigma, M1523-10MG) dissolved in fresh ACSF and injected 250 µL at 20 nL/min at a depth of 500 µm. We recorded the effects of muscimol up to 4 hr post injection. As a control, we injected ACSF-only at the same rate and depth prior to muscimol.

### Traveling wave simulations

We adapted a large-scale spiking network model introduced in previous work ^4^. The model is composed of leaky integrate-and-fire (LIF) neurons, with 80% excitatory and 20% inhibitory, operating in a balanced state ^100^ characteristic of the barrel cortex *in vivo* ^101,102^. Neurons in the model are arranged on a grid, with up to 450,000 neurons distributed over a 4×4 mm^2^ area, with periodic boundary conditions. A large number of synapses per cell (1,000) allows synapses in the model to approximate the scale observed *in vivo* ^103^ (, in contrast to models with smaller numbers of synapses per cell, where recurrent synapses need to be stronger and correlations are overexpressed ^104^. Axonal conduction delays increase linearly with distance between pre- and post-synaptic cells:

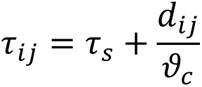

Where *τ_ij_ I*is the delay from neuron *i* to neuron *j*, and *τ_s_* is a fixed delay accounting for synaptic vesicle release, *d_ij_* is the Euclidean distance between neurons *i* and *j*, and *ϑ_c_* is the axonal conduction speed. This network generates self-sustained activity with spontaneous spiking fluctuations consistent with the asynchronous-irregular regime^100,105^. A simulated LFP is calculated from summed excitatory and inhibitory synaptic activity in adjacent, non-overlapping pools of 10 x 10 neurons (corresponding to 67.8 x 67.8 μm^2^)^106^ and is compared to the LFP in the electrophysiological recordings. Simulations were performed using custom software programmed in C (NETSIM; http://mullerlab.github.io). Equations in the model were integrated using the forward Euler method with a time step of 0.1 ms. We then studied the effect of excitatory feedback projections on traveling waves generated by this network. Feedback projections were modelled as external Poisson inputs delivered for a short time window (20 ms) while an individual wave propagates across the network. We focused on feedback delivered only to a local patch of the network, and compared the dynamics in the local network during stimulation to the dynamics before stimulation and in the network outside this local patch. Next, we studied how varying the probability with which the feedback projections target excitatory or inhibitory neurons increases the sparsity of neuronal spiking while the wave propagates across the network.

### Analysis: Data inclusion criteria

For each animal, we recorded 50-300 passive-touch trials. During post-processing, we considered a trial to be “responsive” if a detectable LFP was observable within 70 ms of touch. We defined a detectable LFP as a touch-evoked signal with a maximum at least 3X greater than the pre-stimulus standard deviation (500 ms window prior to touch). All further analyses we performed on these “touch-responsive trials,” which typically consisted of >80% of trials.

### Analysis: Generalized phase

We used the generalized phase (GP) approach to find the instantaneous phase of both the wideband- and narrowband-filtered data (Davis et al., 2020). We filtered all data with a 4^th^ order Butterworth filter (MATLAB’s *filtfilt* function) in the relevant frequency bands and down sampled the LFP to either 2500 Hz or 1000 Hz. To attain the phase of the LFP signal, we used the GP method for calculating the analytic signal. This technique is described in depth elsewhere^5^ and summarized here. This technique is very similar to existing traveling-wave analyses that leverage narrowband filtering ^1,3,13,20^. This approach yields a real-valued, filtered LFP signal, *θ*(*x*, *y*, *z*), and uses a Hilbert transform to calculate the analytic signal *X*:

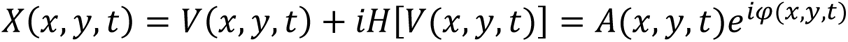

Where *X* is the analytic signal, *V* is the LFP, *H* is the Hilbert transform, *A* is the instantaneous amplitude, *φ* is the instantaneous phase, *x*, *y* are the electrode location and the computation occurs at each timepoint *t*. Typically, the Hilbert transform is calculated using a single-sided Fourier transform (FFT, see MATLAB’s *hilbert* function). In the analytic signal, sinusoids in the LFP are represented in the complex plane as circular contours and the instantaneous phase can then be calculated from the arctangent function (MATLAB’s *angle* function). From the instantaneous phase, the instantaneous frequency can be calculated as the time derivative:

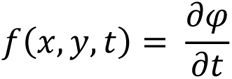

Fast fluctuations in the real LFP signal, *V*(*x*, *y*, *z*), can drive dramatic changes in the instantaneous phase, *φ*(*x*, *y*, *z*), and lead to discontinuities in *f*(*x*, *y*, *z*). Narrowband filtering circumvents these problems, and if discontinuities do occur the most common method of removing these frequencies is to set them to zero (see MATLAB’s *hilbert* function).

The GP approach can be applied to wideband-filtered LFP by solving this discontinuity problem. Like a conventional wave analysis method, the GP approach first calculates the single-sided FFT of the filtered LFP. The GP algorithm then accounts for frequency discontinuities through a shape-preserving interpolation of *φ*(*x*, *y*, *z*), yielding an instantaneous phase that matches the large signal fluctuations at a given point in time, rather than smaller, fast frequencies. The toolbox for performing the computation is publicly available (https://github.com/mullerlab/generalized-phase).

### Analysis: Traveling wave detection with generalized phase

After calculating the instantaneous phase of either the wideband or narrowband LFP with the GP method, we detected traveling waves by analyzing the circular-linear correlation between phase and distance. These methods are outlined in **Figure S2**. First, we detect significant crossings of -π/2 in the phase domain (i.e., positive-rising potentials in the LFP), and label these as “evaluation points” where TWs potentially exist (**Figure S2A**). For these evaluation points to be categorized as a true TW, the phase delays across the grid must follow a linear pattern outward from a source point (i.e. the initiation point of the wave). We found this putative source using functions in the Wave toolbox (Muller et al., 2016) (https://github.com/mullerlab/wave-matlab), which finds the point of maximum divergence of the phase across the grid (**Figure S2B**):

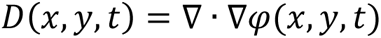

Where *φ*(*x*, *y*, *z*) is the phase across the grid at time point *t*, ∇*φ*(*x*, *y*, *z*) is the phase gradient and ∇·∇*φ*(*x*, *y*, *z*) is the divergence of the phase gradient, *D*(*x*, *y*, *z*). The putative source point, *S*, of the wave is the point of maximum divergence:

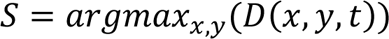

The electrode location at the source point *S* is then considered the site of wave initiation. Moving outwards from this source point, a TW should exhibit a linear increase in phase with distance. A constant phase, or phase that is uncorrelated with distance is not considered a TW. We calculated this correlation using the circular-linear phase vs. distance correlation using the CircStats toolbox (Berens, 2009), which yields a correlation value between 0 and 1 for all potential waves. Waves with a high correlation are considered true TWs (**Figure S2C**). To determine the correlation threshold for TWs, we created a “null distribution” of correlation values by randomly shuffling the electrode locations across the grid. We ran our GP pipeline on this shuffled data to detect evaluation points (i.e., pseudo-TWs), and measured the phase vs. distance correlation values for each point. Iterating through this shuffling process formed a large null distribution low correlation value. We used the 99^th^ percentile of the distribution as a threshold for TW detection in unshuffled LFP (**Figure S2C**). This process was performed for each animal and a unique threshold used for each animal, usually between 0.5-0.6.

The phase vs. distance (i.e., *φ* vs *x*) relationship for each TW yields valuable information. The wave number *k* follows from the spatial derivation:

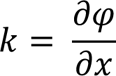

In units of radians/meter. The instantaneous frequency is the time derivative:

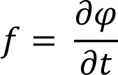

In units of Hz. The instantaneous angular frequency is:

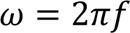

In units of radians/second. Finally, the wave speed can be calculated:

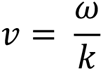

In units of meters/second.

The GP pipeline labels the wave onsets (i.e., -π/2 phase crossings), which are the positive LFP upswings of the wave. To find the amplitude of each wave, we found the LFP amplitude of the next zero-phase crossing point, which labels the LFP peaks.

### Analysis: Phase alignment and phase jitter

Similar to previous studies^5^, we quantified the alignment of the NeuroGrid phase across trials using the standard formulation of the Kuramoto order parameter, which provides a measure of synchrony for coupled oscillators^107^. Running this analysis across all active-touch-evoked LFP yields the average phase alignment on the NeuroGrid and indicates the level of phase synchrony of the recording electrodes (**Figure 3F**). We observed clear local maxima and minima features within the phase alignment data (**Figure 3F**), with small deviations in their time alignment. To quantify this time jitter in Go and No-Go trials, we bootstrapped the data. We randomly chose 100 touch times, calculated the phase alignment across this subset of touches, found the timepoints of distinct maxima and minima for each electrode, then calculated the jitter across electrodes by taking the standard deviation of these timepoints. We repeated this process 1000 times for Go and No-Go trials for each animal, forming a distribution of jitter values (**Figure 3G**).

### Analysis: Late wave detection and amplitude

To sort trials based on the strength of the late wave, we first normalized the single-trial LFP to the mean of the 500 ms pre-stimulus period. We considered the baseline of the recording to be the standard deviation of the pre-stimulus period. We then found the amplitude of the late wave in a window 80-200 ms post-touch. If this amplitude exceeded 3 baseline standard deviations, we considered the trial to have a detectable late wave. Trials under this threshold were categorized as weak late-wave trials. Due to the small variation of the late wave across the grid, we only performed this sorting using one representative electrode chosen by the largest LFP amplitude. We found similar sorting results when using the LFP power, rather than the amplitude.

To measure the average amplitude of the late wave, for each channel we averaged the LFP across all touch-responsive trials in the data. Using a custom MATLAB GUI, we plotted these waveforms, annotated the approximate onset of the late spike, then ran an algorithm that automatically detected the late wave onset (the trough) and the late wave peak (the maximum 100-300 ms after touch). The late wave amplitude for the channel is then computed as the trough-to-peak amplitude (see **Figure S8B**).

### Analysis: Removing two-photon scanning artifacts on NeuroGrids

Resonant laser scanning during two-photon imaging led to mild noise artifacts on the NeuroGrid data (see **Figure S1D-E**). To remove these artifacts, we filtered the LFP data with multiple 6th order Butterworth notch filters at frequencies of 30 Hz and its harmonics up to 360 Hz. This approach removed the majority of the noise power while preserving the overall LFP (**Figure S1E**).

### Analysis: two-photon calcium imaging segmentation

Each two-photon imaging trial generated a 32-bit TIFF stack. For each animal, we concatenated all stacks using ImageJ, denoised with a Kalman stack filter plugin (using default parameters), and converted the concatenated files to 8-bit format. We used a CaImAn pipeline to segment regions of interest (ROIs) and extract calcium traces from two-photon imaging data^108^. Due to our large data sets, we used the “patches” functionality, which analyzes smaller patches of the full FOV in parallel. This pipeline includes motion correction using NormCorr, segmentation using constrained non-negative matrix factorization, neuropil subtraction, and deconvolution to extract calcium events. Lastly, features were refined through a convolved neural network classifier. We determined two parameters that were key to dendritic segmentation. We lowered the classifier threshold (‘cnn_thr’) from 0.4 to 0.3, and the NMF converging coefficient from 0.8 to 0.5. We found that the “patches” functionality labeled neurons well, but also provided a number of extraneous ROIs. We manually curated the segmentation post-hoc and removed these erroneous ROIs with a custom MATLAB GUI.

We classified cells as touch-responsive if the averaged touch-evoked ΔF/F transient showed a maximum value 3X larger that the baseline standard deviation within 2 seconds of touch. To calculate the touch-evoked onsets of each cell on a single-trial basis, we first calculated an event threshold for each cell using the 99^th^ percentile of all events in the deconvoluted data. Events that exceeded this threshold were considered true calcium events. We then determined if an event occurred within a 1 s window after each touch stimulus. If an event occurred, the largest calcium event within the window in the deconvoluted data is maximum slope of the ΔF/F upswing (**Figure 7G**). We found the onset of the event by finding the preceding inflection point in the deconvoluted traces (**Figure 7F**).

### Analysis: Calcium data vectorization, reactivation index, and ensemble detection

Our ensemble analysis pipeline is outlined in **Figure S5** and follows previous reports ^53,109^. Changes in fluorescence and convolved spiking activity were filtered by omitting events below 2.5 standard deviations of the noise floor. This ensured robust spiking analysis for neuronal population networks. We first generated a N x T calcium activity matrix by plotting the mean centered fluorescence values for all detected cells, where N denotes the number of neurons and T denotes the total number of frames for each video:

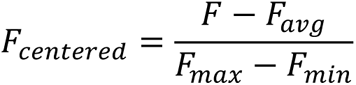

We then used a coactivity threshold to transform this calcium activity matrix into N x T binary spike matrix (**Figure S5A**). Each row (N) represents the spiking events of a neuron. The threshold was determined by shuffling the data (see below) and essentially removes solitary calcium events, focusing our analyses on periods of coactivity. We then computed a vectorized form of the binary matrix by binning frames every 300 ms and summing the calcium events across time in each bin (**Figure S5A**). This summation produced a matrix of time vectors, *t_i_* = (*N_i_*, *T_i_*), where *N_i_* is the summed event spikes of time bin *T_i_*. This vector series represents temporal changes of grouped neurons.

This vectorized form of calcium activity is advantageous, since now vector-based methods can be used to for correlating changes of network states between vector pairs *t_i_* and *t_j_* using a cosine-similarity index (**Figure S5B**):

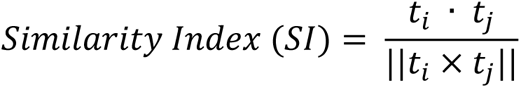

This yielded a non-negative value corresponding to the cosine angle between any vector pair in vector space (**Figure S5B**). A value near 1 corresponds to vectors with nearly identical activity patterns within the binned time frame. Running this calculation for all vectors pairs yields a *T_i_xT_i_* “similarity map” that indicates periods of similar patterns of activity across the FOV (**Figure S5B**). We then further refine this similarity map by removing all vectors that do not show a significant similarity with any vector pairs (threshold determined by shuffling the data), which we refer to as the reactivation index as this metric indicates the prevalence of similar reactivation patterns across the imaging trial.

The similarity map for an animal highlights periods of similar activity patterns during the recording. To identify significant patterns in the matrix, we found the principal components of the high-dimensional data using singular-value decomposition (SVD). SVD allows for the identification of components that contribute the most to a given network state. The SVD of a matrix can be represented as:

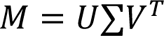

Where, *M* is the similarity matrix, *U* and *V* are the orthonormal bases, and ∑ are the eigenvalues of the principal components in matrix *M*. In our case, because *M* is symmetric (see **Figure S5B**), this reduces to:

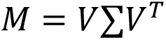

We found the eigenvalues that captured the majority of the data variance by comparing the SVD to a shuffled data set. This typically left 3-10% of eigenvalues per animal that capture > 90% of the data variance. Activity vectors are then grouped and spatially projected using these eigenvalues and reproduce the main activity patterns of the original data to a high degree. These eigenstates are used as candidates for neuronal ensembles.

Importantly, each eigenstate (i.e., activity state) is a vector composed of the activity of all neurons within the FOV. However, only a subset of neurons within this vector are co-active and form the majority of the activity within the eigenstate. These active cells compose the neuronal ensemble. To find these active cells, we used a SØrenson-Dice correlation (SDC) to quantify synchronous firing events of individual neurons (**Figure S5C)**. Unique neuronal pairs identified through the binary matrix (N x T) are linearized and filtered through a function:

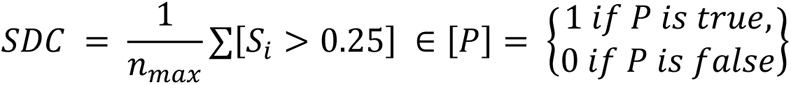

The resulting correlation matrix represents event pairing of identified neurons throughout the video and yields critical factors such as node strength (i.e., a measure of activity), functional connectivity strength between neuron pairs, and numbers of functional connections (**Figure S5C**). These attributes both identify the neuronal ensemble that composes the activity state and also yields network functional connectivity metrics. The proportion of neurons within an ensemble that were coactive above shuffled chance was defined as the ensemble reactivation.

To set ensemble detection thresholds, multiple data shuffling procedures were performed. Initial spikes per neuron in the binary matrix N x T were temporally shuffled across each frame of the time series. An event shuffle is further introduced between two temporally shuffled ROIs to produce a spatiotemporal shuffled binary matrix. This binary surrogate is repeatedly shuffled. Shuffling was done by generating a random permutation equal to the length of the neurons and frames. Spikes were then indexed to each random number generated. This method randomizes the network state rendering the population of the spiking as stochastic events.

### Analysis: Single-unit recordings

We pre-processed data by bandpass filtering with a 4^th^-order Butterworth filter between 300-3000 Hz (MATLAB’s *filtfilt*). After filtering, we subtracted the median signal across all channels from each recording site (i.e., common-median referencing) to remove shared high-frequency noise^110^. We used KiloSort2 for spike sorting silicon probe recordings (https://github.com/MouseLand/Kilosort/releases/tag/v2.0) and manually curated the clustered data using Phy2 (https://github.com/cortex-lab/phy/) ^111^

### Analysis: Current-source density

We used current-source density (CSD) analysis to determine current sources and sink across a linear silicon probe during LFP recordings ^112^. The CSD is calculated from the second spatial derivative the LFP transients:

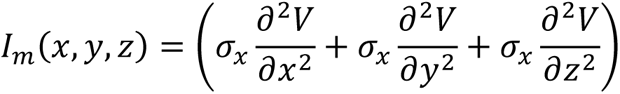

Where *I_m_* is the CSD, *V* is the LFP, and *σ* are the spatial conductivities of the extracellular space. When assuming laminar uniformity and constant conductivity, the derivatives in the x and y directions becomes zero and this calculation can be approximated as:

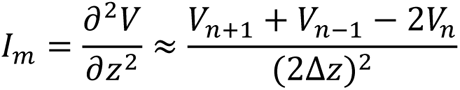

Where *I_m_* is the CSD, *V_n_* is the LFP on the *n*th electrode, and Δ*z* is the electrode spacing (Freeman and Nicholson, 1975). For our calculations, we used n = 2 and added two Vaknin electrodes ^113^ at the top and bottom of the silicon probe to estimate the CSD across the full shank length.

### Analysis: Spectral analysis of grid and shank recordings

To calculate LFP spectrograms from NeuroGrid recordings (**Figure 4C**) and shank recordings (**Figure S4C**), we chose a representative electrode and used a continuous wavelet transformation (MATLAB’s *cwt* function) to calculate the single-trial spectrogram between 0.1 and 100 Hz. The single-trial spectrograms for all touch-responsive trials were then averaged to generate the average spectrogram.

To calculate the beta/gamma and theta/gamma power ratios from NeuroGrid recordings (**Figure 4D**), we first calculated the average spectrogram from a representative channel as outlined above. We then summed all the data points in the spectrogram in the respective theta, beta, and gamma bands to get the “total power” in the early (0-60 ms window) and late (100-200 ms window). The ratios of the total power yielded the relative beta/gamma and theta/gamma power during the early and late wave periods.

To calculate the relative power profile in laminar shank recordings (**Figure S4B**), we mirrored previously reported methods^114^. Because the UCLA silicon probe recording electrodes are organized into a hexagonal format and offset in the x-axis, we limited our analyses to the linearly aligned electrodes on either the left- or right-hand side of the shank. First, we focused on a time window between spanning - 100 ms to +200 ms around the touch stimulus. For each laminar electrode, we then calculated the average power spectral density across trials (MATLAB’s *pwelch*). We then concatenated the average PSD vectors for every laminar electrode into a *f x N* matrix where *f* is the PSD frequency and *N* is the number of channels organized from superficial electrodes to deep electrodes. The relative power at each frequency point is calculated by normalizing each column of the matrix to its maximum value. This process can be summarized with the following equation:

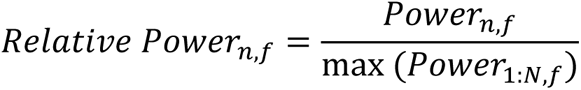

Where *n* is the electrode channel, *N* is the total number of electrodes and *f* is the frequency (Mendoza-Halliday et al., 2023).

### Statistical analyses

All statistical testing was performed in MATLAB. For boxplots, the upper/lower edges of the box represent the upper/lower quartiles of the data. The line within the box represents the median of the data. Data points that fall outside the boxplot upper/lower “whiskers” are considered outliers. We tested data for normality using a Kolmogorov–Smirnov test. In all data sets within this study, we determined at least one group per comparison to be non-normally distributed and therefore we used only non-parametric statistical tests. Statistical details can be found within the legends of each figure. In short, we used a ranked-sum Wilcoxon test (otherwise known as a Mann-Whitney U-test) to compare unpaired, non-parametric data between two groups. We used a signed-rank Wilcoxon test to compare paired, non-parametric data between two groups. We used a Kruskal-Wallis test to compare unpaired, non-parametric data between three or more groups and followed this with a *post-hoc* Dunn-Sidak test to correct for multiple comparisons. Finally, we used a Friedman’s test to compare paired, non-parametric data between three or more groups and followed this with a *post-hoc* Dunn-Sidak test to correct for multiple comparisons. In some cases, we used bootstrapping to create data distributions to compare two conditions. In **Figure 3G**, for each animal we randomly chose 100 touch times, calculated the mean touch-evoked LFP, calculated the phase alignment across the grid, then calculated the phase jitter across the grid. We repeated this process 1000 times for each animal to create the Go and No-Go data distributions. In **Figure 7G**, we randomly chose 100 dendrites (with resampling), and calculated the standard deviation of ΔF/F onset times. We repeated this process 10,000 times for the late-wave and weak-late-wave conditions to form a distribution standard-deviation values for each condition to compare with a ranked-sum Wilcoxon test. In **Figure S7C**, we randomly selected 20 trials with and without the late wave, calculated the mean LFP, calculated the CSD, then measured the magnitude of the late L2/3 current sink. We repeated this process 50 times for each animal to form distributions of the current sign magnitudes with and without the late wave. Reported n-values and p-values can be found within the test and legends for each data comparison.

