## Supporting Information for "A Translaminar Spacetime Code Supports Touch-Evoked Traveling Waves"

#### TITLE

#### Authors and Affiliations

Daniel L. Gonzales[1], Hammad F. Khan[1], Hayagreev V.S. Keri[1-2], Saumitra Yadav[1], Christopher Steward[6], Lyle E. Muller[4-5], Scott R. Pluta[2-3], and Krishna Jayant[1,3,7]

[1]Weldon School of Biomedical Engineering, Purdue University, West Lafayette, IN 47907, USA

[2]Department of Biological Sciences, Purdue University, West Lafayette, IN 47907, USA

[3]Purdue Institute for Integrative Neuroscience, Purdue University, West Lafayette, IN 47907, USA

[4]Department of Applied Mathematics, Western University, London, ON, Canada

[5]Brain and Mind Institute, Western University, London, ON, Canada

[6]Department of Computer Science, Western University, London, ON, Canada

[7]Corresponding author and lead contact

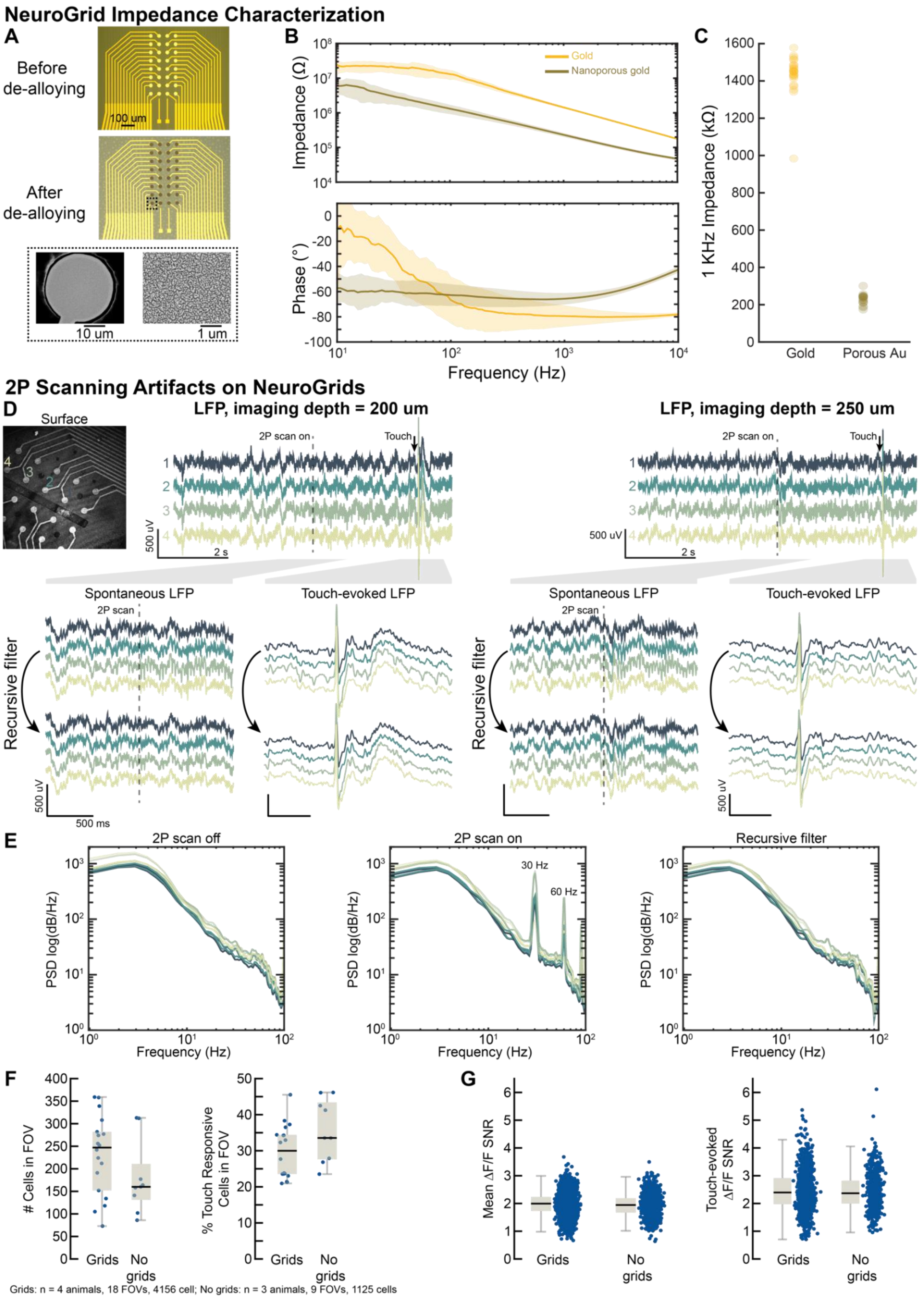

NeuroGrid Impedance Characterization
A
Before de-alloying
100 um
After de-alloying
10 um
1 um
B
Gold
Nanoporous gold
Impedance (Ω)
Phase (°)
Frequency (Hz)
C
1 KHz Impedance (kΩ)
Gold
Porous Au
2P Scanning Artifacts on NeuroGrids
D
Surface
LFP, imaging depth = 200 um
LFP, imaging depth = 250 um
2P scan on
Touch
500 uV
2 s
Spontaneous LFP
Touch-evoked LFP
2P scan
Recursive filter
500 ms
E
2P scan off
2P scan on
Recursive filter
30 Hz
60 Hz
PSD log(dB/Hz)
Frequency (Hz)
F
### Cells in FOV
% Touch Responsive Cells in FOV
Grids
No grids
Grids: n = 4 animals, 18 FOVs, 4156 cell; No grids: n = 3 animals, 9 FOVs, 1125 cells
G
Mean ΔF/F SNR
Touch-evoked ΔF/F SNR


**Figure S1**

**Figure S1. NeuroGrid electrochemical impedance and imaging characterization.**

(A) Optical micrographs of an on-chip NeuroGrid before and after the de-alloying process in nitric acid. Insets are scanning-electron micrographs of one recording pad, which shows the porosity.

(B) Impedance and phase comparison of recording sites composed of only gold and gold+nanoporous gold.

(C) 1 KHz impedance comparison of gold vs nanoporous gold recording sites.

(D) (Top) Micrograph shows NeuroGrid orientation on the brain surface. Selected channels are labeled. LFP traces show recordings from selected recording sites at imaging depths of 200 and 250 μm. Two-photon laser scanning turns on halfway through the traces (dashed line), and a touch occurs near the end. (Bottom) LFP traces when the two-photon scanning turns on and when touch occurs. The arrow indicates the same traces after applying a recursive notch filter to remove scanning noise. Scale bars: 500 μV and 500 ms.

(E) Power spectrum during periods with no imaging (left), during two-photon imaging (center), and two-photon imaging data after the recursive filter is applied (right). Power spectrums were calculated during spontaneous LFP activity.

(F) Total number of cells and touch-responsive cells in wS1 in the FOV with and without simultaneous surface grid recordings.

(G) Comparison of the $\Delta F/F$ SNR for all calcium events (left) and touch-evoked calcium events (right) during recordings with and without the NeuroGrid.

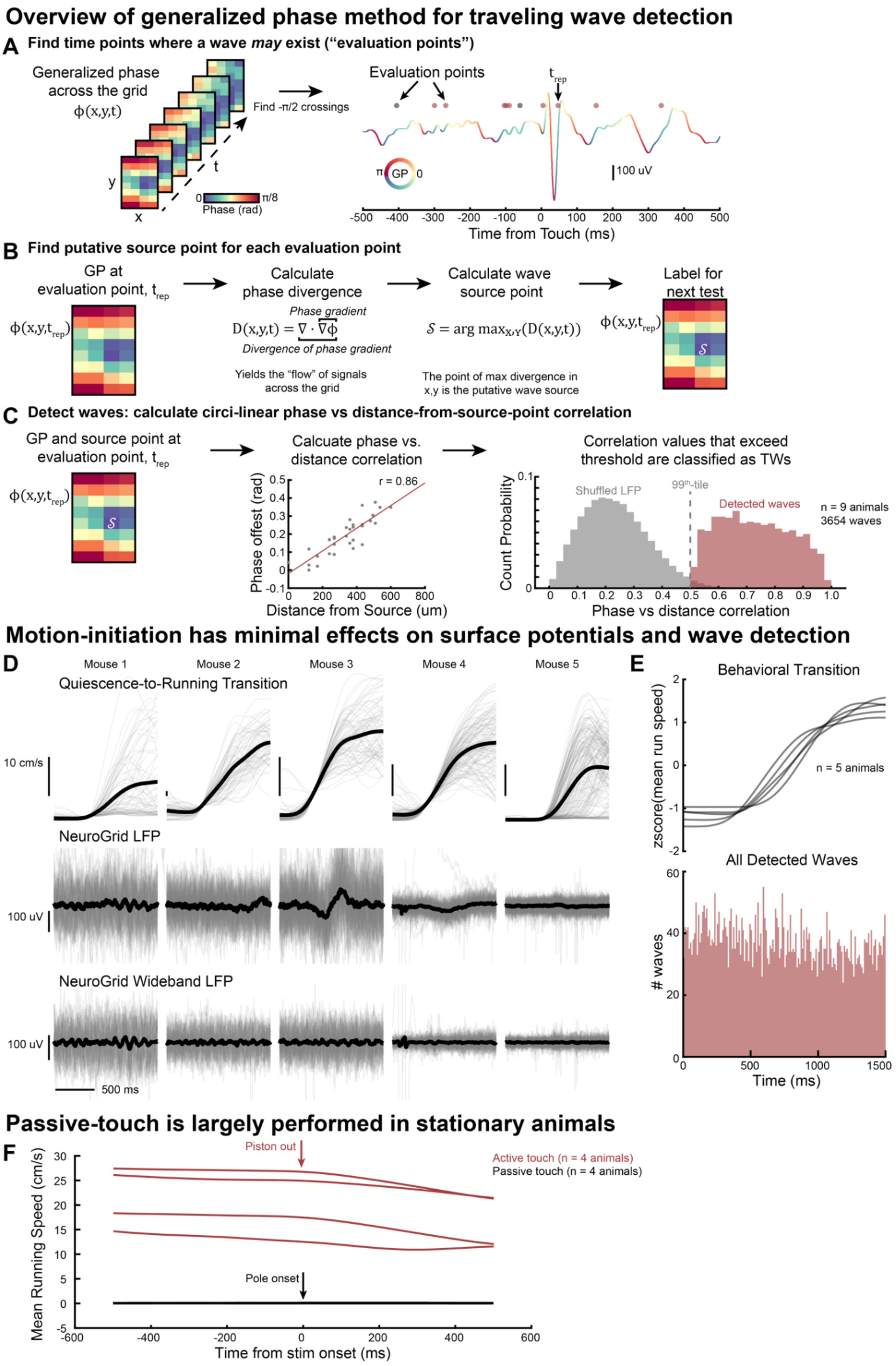
Overview of generalized phase method for traveling wave detection
A Find time points where a wave may exist ("evaluation points")
Generalized phase across the grid
ϕ(x,y,t)
Find -π/2 crossings
Phase (rad)
0
π/8
Evaluation points
t_rep
GP
100 uV
Time from Touch (ms)
B Find putative source point for each evaluation point
GP at evaluation point, t_rep
ϕ(x,y,t_rep)
Calculate phase divergence
Phase gradient
D(x,y,t) = ∇ · ∇ϕ
Divergence of phase gradient
Yields the "flow" of signals across the grid
Calculate wave source point
S = arg max_x,y(D(x,y,t))
The point of max divergence in x,y is the putative wave source
Label for next test
C Detect waves: calculate circi-linear phase vs distance-from-source-point correlation
GP and source point at evaluation point, t_rep
Calcuate phase vs. distance correlation
r = 0.86
Phase offest (rad)
Distance from Source (um)
Correlation values that exceed threshold are classified as TWs
Shuffled LFP
99th-tile
Detected waves
n = 9 animals
3654 waves
Count Probability
Phase vs distance correlation
Motion-initiation has minimal effects on surface potentials and wave detection
D
Mouse 1
Mouse 2
Mouse 3
Mouse 4
Mouse 5
Quiescence-to-Running Transition
10 cm/s
NeuroGrid LFP
100 uV
NeuroGrid Wideband LFP
500 ms
E
Behavioral Transition
zscore(mean run speed)
n = 5 animals
All Detected Waves
### waves
Time (ms)
Passive-touch is largely performed in stationary animals
F
Piston out
Active touch (n = 4 animals)
Passive touch (n = 4 animals)
Pole onset
Mean Running Speed (cm/s)
Time from stim onset (ms)

Figure S2

**Figure S2. Overview of generalized phase pipeline and locomotion effects on NeuroGrid LFP**

(A) Workflow for finding points-of-interest for potential TW detection. (Left) Representative time points of generalized phase across the grid calculated from the LFP. (Right) We detected all $-\pi/2$ phase crossings in the touch trial and label these as "evaluation points" for more testing to determine if a TW exists.

(B) For each evaluation point, we determined the putative source point by calculating the divergence of the phase gradient across the grid.

(C) For each evaluation point, we then calculated the circular-linear phase vs. distance-from-source-point correlation. Points with strong correlation values were then determined to be TWs. Significant correlation values were found by first constructing a "null distribution" of correlation values for each animal. We did this by shuffling the LFP location across the grid, running GP analysis, and calculating the correlation value for all pseudo-evaluation points (50 shuffle iterations were performed for each trial for each animal). The $99^{th}$ percentile of the null distribution was used as a threshold for traveling wave detection in unshuffled data. In panel (A), black evaluation points had correlation values below the threshold.

(D) We considered whether the late wave could be confounded by delivering a passive touch to quiescent animals that drives a behavioral state transition to locomotion. We analyzed animal locomotion and detected spontaneous behavioral transitions from quiescence to running in the absence of touch. For these behavioral transitions, we aligned the full LFP spectrum (middle traces) and wideband filtered LFP used for traveling wave analysis (bottom traces). We observed no consistent influence of behavioral state transitions on the NeuroGrid LFP.

(E) For the behavioral transitions in the absence of touch, we also performed traveling wave detection. We detected spontaneous waves in the LFP, but they had no observable consistency with the onset of locomotion.

(F) Average running speed across all animal trials during the active and passive touch paradigms.

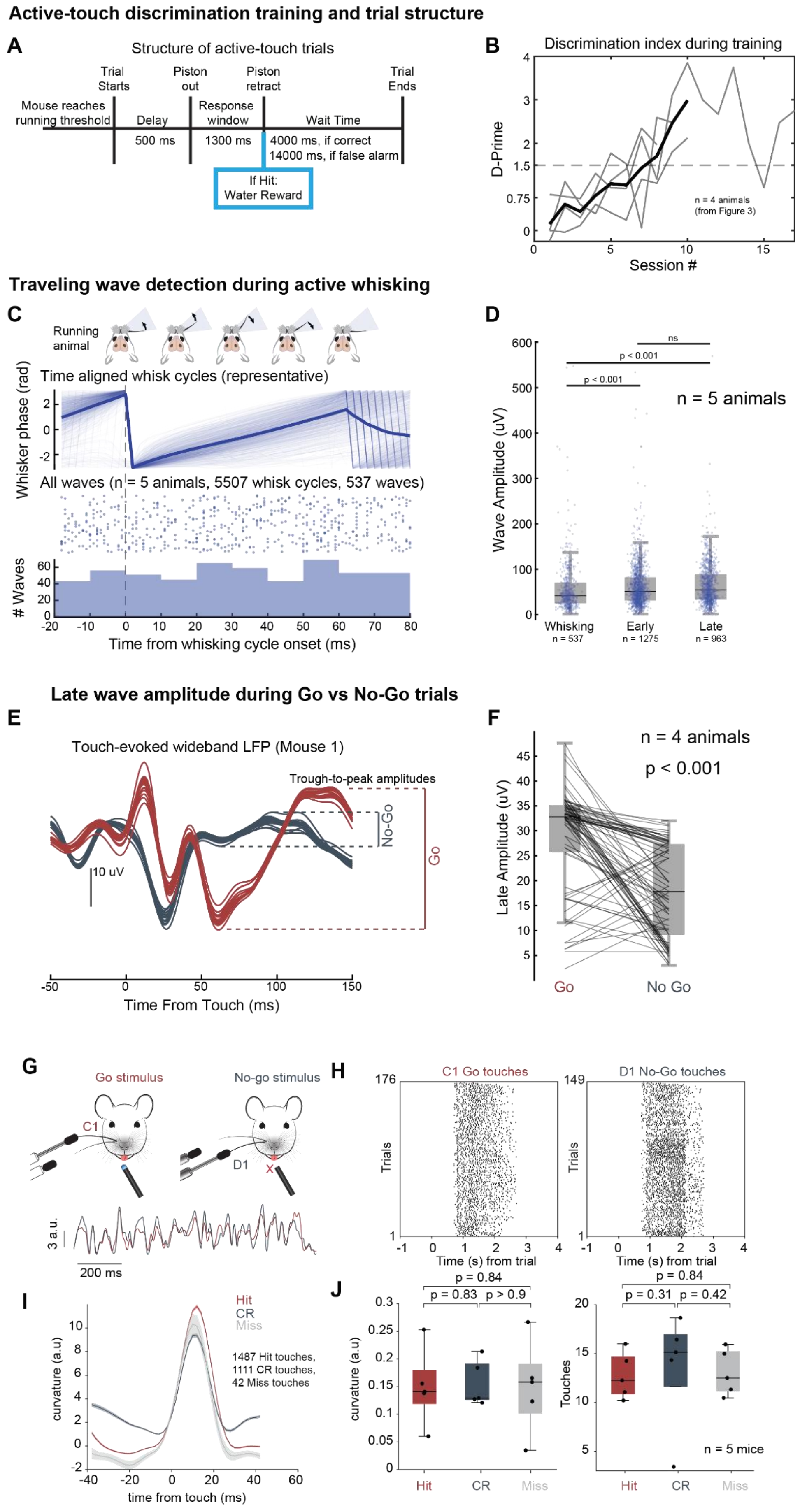
Active-touch discrimination training and trial structure
A
Structure of active-touch trials
Trial Starts
Piston out
Piston retract
Trial Ends
Mouse reaches running threshold
Delay
Response window
Wait Time
500 ms
1300 ms
4000 ms, if correct
14000 ms, if false alarm
If Hit: Water Reward
B
Discrimination index during training
D-Prime
Session #
n = 4 animals (from Figure 3)
Traveling wave detection during active whisking
C
Running animal
Time aligned whisk cycles (representative)
Whisker phase (rad)
All waves (n = 5 animals, 5507 whisk cycles, 537 waves)
### Waves
Time from whisking cycle onset (ms)
D
Wave Amplitude (uV)
n = 5 animals
ns
p < 0.001
p < 0.001
Whisking
n = 537
Early
n = 1275
Late
n = 963
Late wave amplitude during Go vs No-Go trials
E
Touch-evoked wideband LFP (Mouse 1)
Trough-to-peak amplitudes
No-Go
Go
10 uV
Time From Touch (ms)
F
n = 4 animals
p < 0.001
Late Amplitude (uV)
Go
No Go
G
Go stimulus
No-go stimulus
C1
D1
3 a.u.
200 ms
H
C1 Go touches
D1 No-Go touches
Trials
Time (s) from trial
I
Hit
CR
Miss
1487 Hit touches,
1111 CR touches,
42 Miss touches
curvature (a.u)
time from touch (ms)
J
p = 0.84
p = 0.83
p > 0.9
p = 0.84
p = 0.31
p = 0.42
curvature (a.u)
Touches
n = 5 mice
Hit
CR
Miss

**Figure S3.**

**Figure S3. Overview of two-whisker, active-touch discrimination behavior.**

(A) Details of the single-trial structures conducted during the active-touch paradigm.

(B) Discrimination index for the two-whisker operant behavior during training sessions. We considered a discrimination index of 1.5 as expert behavior.

(C) We considered that whisking itself may evoke wS1 traveling waves. We detected waves during free whisking with no touch-related objects to test this. (Top) Whisking phase. The schematic shows different whisker positions and their relationship to the whisking phase. The plot shows time-aligned whisking phases for a representative animal. We limited our analyses to whisking cycles that lasted a minimum of 60 ms. (Middle) Raster plot of all detected traveling wave onsets in the wideband LFP across all animals. (Bottom) Histogram of wave onset times. There is no clear correlation between the whisking phase and traveling wave onset (n = 5 animals, 5507 total whisking cycles, 537 total detected waves).

(D) Traveling wave amplitude for free whisking, touch-evoked early waves (0-50 ms post touch), and touch-evoked late waves (50-100 ms post touch). Touch-evoked amplitudes are significantly stronger (n = 5 animals, 537 whisking-only waves, 1275 early waves, 963 late waves, p-values determined with a Kruskal-Wallis test with a *post-hoc* Dunn-Sidak test).

(E) Representative active-touch evoked NeuroGrid LFP for Go (C1 touch) and No-Go (D1 touch) trials. The arrow indicates the trough-to-peak region of the late wave that was quantified.

(F) The late wave trough-to-peak amplitude quantified across all electrodes was significantly higher for Go trials, indicating reward reinforcement enhances the late wave (n = 4 animals, p-value determined with a signed-rank Wilcoxon test).

(G) (Top) Schematic of active behavior paradigm involving two-whisker discrimination. (Bottom) Trial average whisking response for the representative animal.

(H) A representative touch raster across one recording session for Go and No-Go trials.

(I) Whisker variance between Hit and Correct reject trials (n = 325 trials).

(J) (Left) No difference was observed in the variation of whisker movement across trials and between task conditions (rank-sum Wilcoxon test, n = 5 mice). (Right) No difference was observed in the number of sampled touches between task conditions (rank-sum Wilcoxon test, n = 5 mice)

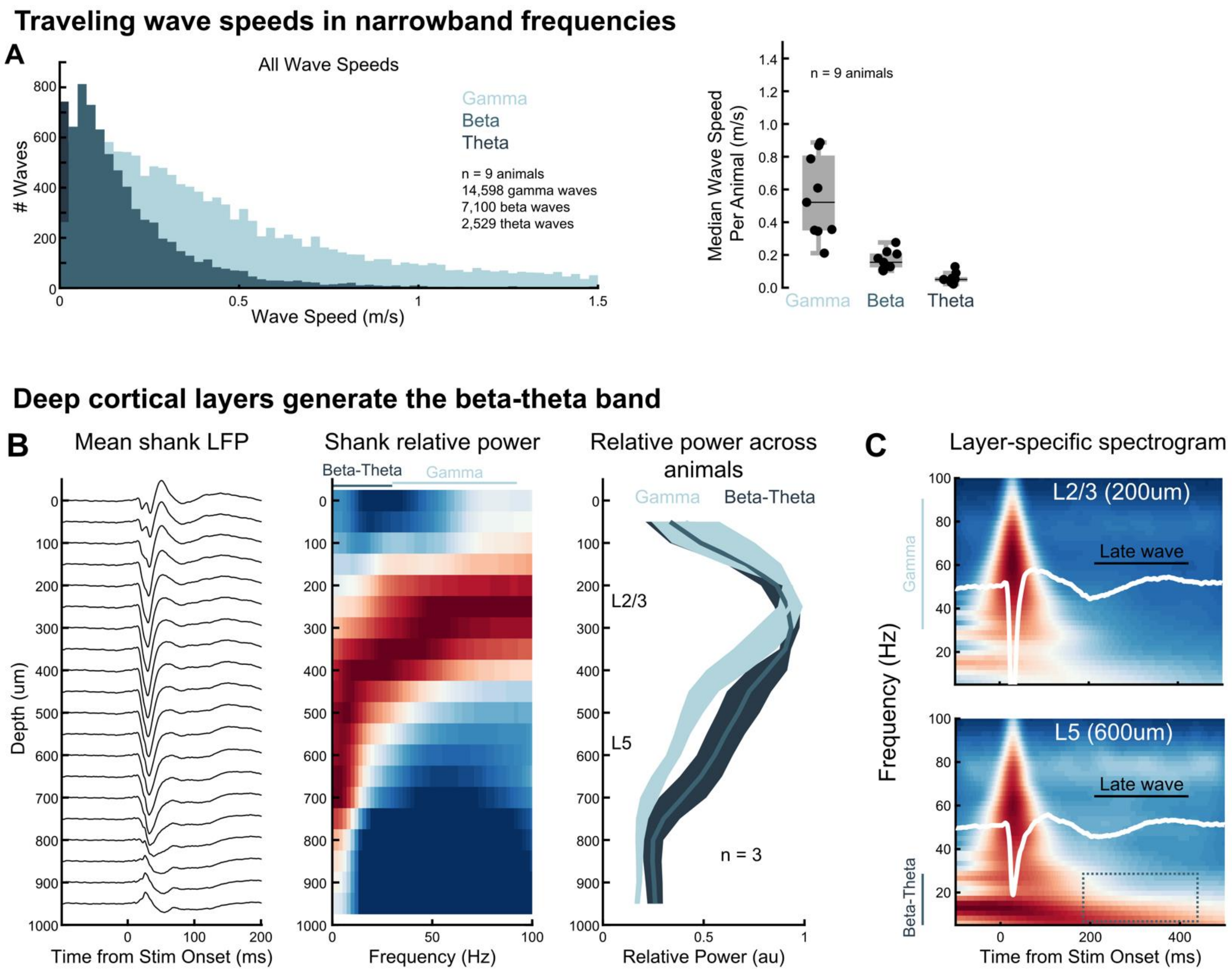


**Figure S4.**

**Figure S4. Narrowband frequency analysis of wS1 traveling waves with NeuroGrids and silicon probes.**

(A) (Left) Speeds for all detected traveling waves on the NeuroGrid in wS1 in the gamma, beta, and theta bands (n = 9 animals, 14,589 gamma waves, 7,100 beta waves, 2,529 theta waves). (Right) Median wave speed per animal for each frequency band.

(B) (Left) Mean touch-evoked LFP on a wS1 silicon probe for a representative animal. (Center) Representative relative power in each frequency band as a function of cortical depth. This laminar profile shows that the infragranular layers (>400 µm) have enhanced beta-theta power (4-30 Hz), and the supragranular layers are dominated by the gamma band (30-90 Hz). (Right) Relative laminar power for the beta-theta and gamma bands across animals (n = 3). Again, the infragranular layers—and most notably L5—are primarily driven by the beta-theta band. Shaded error is the standard error of the mean.

(C) L2/3 and L5 touch-evoked spectrogram for a representative animal. The late wave appears ~200 ms after touch and most clearly correlates with delayed beta-theta coupling in L5.

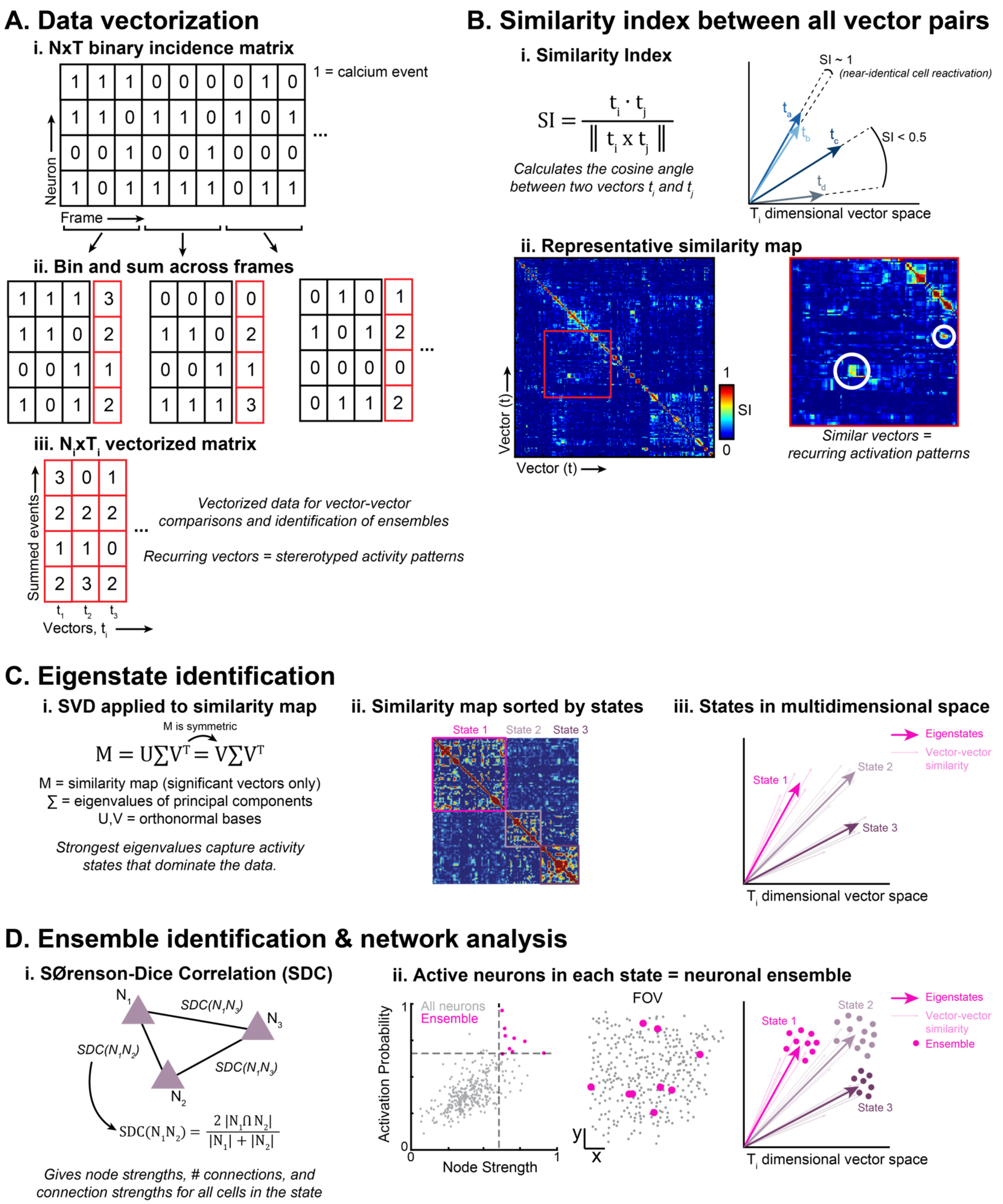
A. Data vectorization
i. NxT binary incidence matrix
1 = calcium event
Neuron
Frame
ii. Bin and sum across frames
iii. $N_i x T_i$ vectorized matrix
Summed events
Vectors, $t_i$
Vectorized data for vector-vector comparisons and identification of ensembles
Recurring vectors = sterertotyped activity patterns
B. Similarity index between all vector pairs
i. Similarity Index
$SI = \frac{t_i \cdot t_j}{\| t_i \times t_j \|}$
Calculates the cosine angle between two vectors $t_i$ and $t_j$
SI ~ 1
(near-identical cell reactivation)
SI < 0.5
$T_i$ dimensional vector space
ii. Representative similarity map
Vector (t)
SI
Similar vectors = recurring activation patterns
C. Eigenstate identification
i. SVD applied to similarity map
M is symmetric
$M = U\Sigma V^T = V\Sigma V^T$
M = similarity map (significant vectors only)
∑ = eigenvalues of principal components
U,V = orthonormal bases
Strongest eigenvalues capture activity states that dominate the data.
ii. Similarity map sorted by states
State 1
State 2
State 3
iii. States in multidimensional space
Eigenstates
Vector-vector similarity
$T_i$ dimensional vector space
D. Ensemble identification & network analysis
i. SØrenson-Dice Correlation (SDC)
$SDC(N_1N_2) = \frac{2\,|N_1 \cap N_2|}{|N_1| + |N_2|}$
Gives node strengths, # connections, and connection strengths for all cells in the state
ii. Active neurons in each state = neuronal ensemble
All neurons
Ensemble
Activation Probability
Node Strength
FOV
Ensemble
$T_i$ dimensional vector space

Figure S5

**Figure S5. Ensemble detection and network connectivity analysis pipeline.**

(A) Calcium event binning and vectorization procedure. With this technique, we can use vector-based calculations to determine common patterns of reactivation across the cortical landscape.

(B) Dot-product vector multiplication for similarity index calculation. This allows for comparing all vectors in the data set. Vectors with a high similarity will have nearly identical cellular activation patterns.

(C) SVD procedure for identifying activity states that dominate the calcium data.

(D) SDC calculations provide detailed network connectivity metrics and the cells that primarily compose the eigenstates.

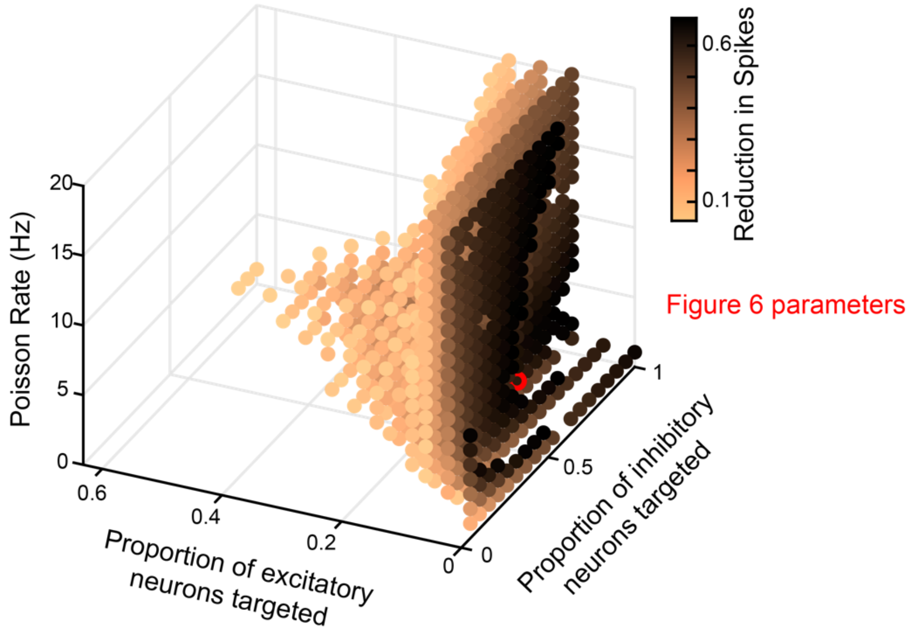


**Figure S6.**

**Figure S6. Feedback simulation with varying E-I ratios.**

Feedback increases the sparsity of traveling waves across a broad range of ratios and firing rates. Plotted are results from simulations of the sparse-wave network, with different proportions of excitatory neurons in the patch targeted by feedback projections, proportion of inhibitory neurons targeted, and Poisson rate for feedback. Each dot plotted in the figure represents a simulation where sparsity was increased by feedback by approximately 75% or higher (measured as the percentage decrease in spiking activity caused by the feedback compared to the without-feedback control). These results demonstrate that, for a broad range of values, feedback that predominantly targets inhibitory neurons can substantially increase the sparsity of waves traveling over the spiking network model.

NeuroGrids + silicon probe confirms the late wave is a superficial current sink

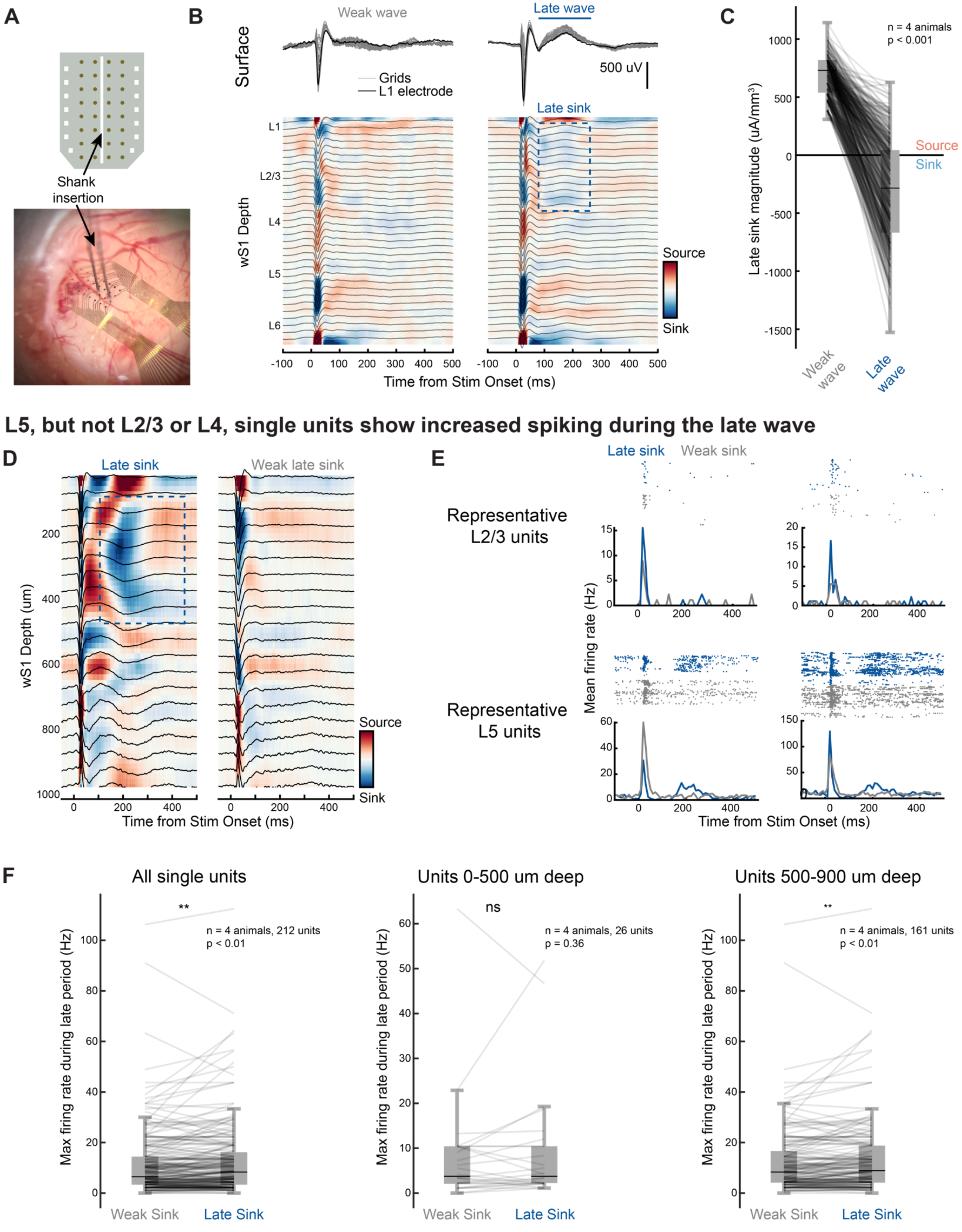


Figure S7.

**Figure S7. The late wave is a supragranular current sink with non-somatic origins.**

(A) Schematic and optical micrograph demonstrating how thin silicon probes are inserted through a small through-hole in the NeuroGrid.

(B) Representative simultaneous surface and depth recording. We sorted NeuroGrid trials based on the strength of the late wave and performed CSD analyses on the corresponding laminar recordings. (Top) Average LFP across all NeuroGrid channels for late-wave and weak-late-wave trials (gray). Also shown is the mean LFP for the superficial shank electrode (black). (Bottom) Average shank LFP for each group and the corresponding CSD from the average waveforms. The late wave correlates with a clear, delayed L2/3 current sink.

(C) Average late sink magnitude during late-wave and weak-late-wave trials. The late sink strengthens with a strong late wave. ($n = 4$ animals, p-value determined with a signed-rank Wilcoxon test following bootstrapping of LFP single trials).

(D-F) Silicon probe recordings with no surface grids for single-unit analyses during the late sink.

(D) Mean touch-evoked LFP across the laminar probe for a representative animal and corresponding CSD. Trials were sorted by the strength of the late L2/3 sink.

(E) Representative single units in L2/3 (200 and 250 µm depth, respectively) and L5 (600 and 650 µm depth, respectively). Both a raster plot of spike times for individual trials and the average firing rate across trials are shown for each unit. Trials were sorted based on the strength of the L2/3 late sink (strong late sink in blue and weak late sink in gray). Notably, L5 neurons show an additional bump in their firing rates during the late sink period (100-300 ms), while L2/3 neurons do not.

(F) We compared strong- vs. weak-late-sink firing rates across all detected single units (left), supragranular single units (center), and infragranular single units (right) ($n = 4$ animals). Across all neurons, we observed an increase in the maximum firing rate during the late period (100-300 ms) in trials with a prominent late sink (left, $n = 212$ units, $p < 0.01$). This increase was not observed in units with a depth of less than 500 µm (center, 26 units, $p = 0.36$). This effect was restricted to infragranular single units with a depth greater than 500 µm (right, $n = 161$ units, $p < 0.01$). A signed-rank Wilcoxon test was used for all comparisons. These results indicate that the superficial late sink is not driven by an increase in superficial-layer spiking. However, increased spiking in L5 neurons during the late sink suggests that L5 spiking drives apical dendrite electrogenesis and the delayed current sink.

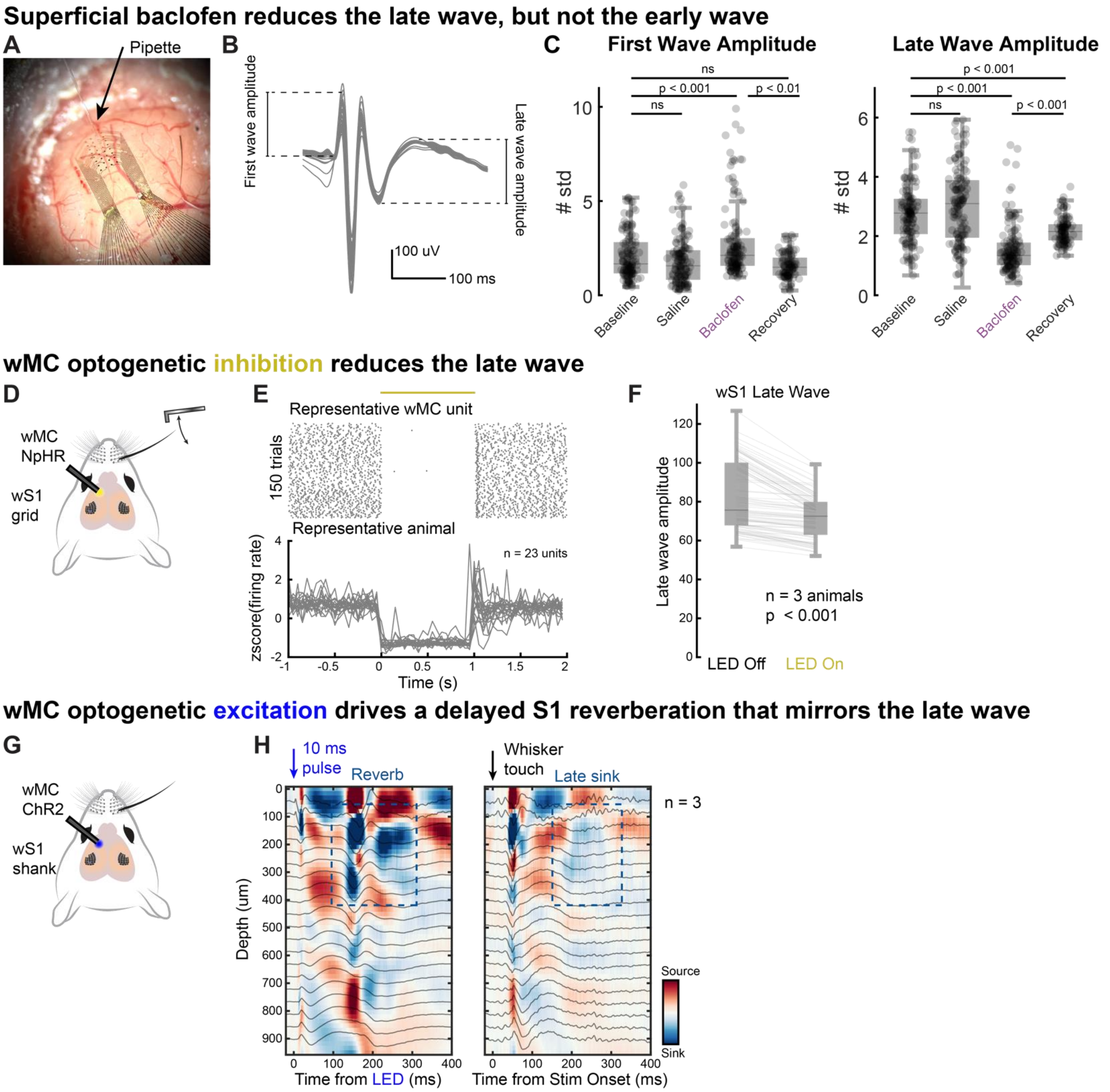
Superficial baclofen reduces the late wave, but not the early wave
A
Pipette
B
First wave amplitude
Late wave amplitude
100 uV
100 ms
C
First Wave Amplitude
Late Wave Amplitude
ns
p < 0.001
p < 0.01
p < 0.001
### std
Baseline
Saline
Baclofen
Recovery
wMC optogenetic inhibition reduces the late wave
D
wMC
NpHR
wS1
grid
E
Representative wMC unit
150 trials
Representative animal
n = 23 units
zscore(firing rate)
Time (s)
F
wS1 Late Wave
Late wave amplitude
n = 3 animals
p < 0.001
LED Off
LED On
wMC optogenetic excitation drives a delayed S1 reverberation that mirrors the late wave
G
wMC
ChR2
wS1
shank
H
10 ms
pulse
Reverb
Whisker
touch
Late sink
n = 3
Depth (um)
Source
Sink
Time from LED (ms)
Time from Stim Onset (ms)

**Figure S8.**

**Figure S8. Extended results from wS1 baclofen injections and M1 optogenetics**

(A) Image showing a puff micropipette inserted into a through-hole for pharmacology directly under the NeuroGrids.

(B) Schematic indicating how the first and late wave amplitudes were calculated from the touch-evoked LFP.

(C) The amplitude of the first wave and late wave during baseline recordings (no injections), saline injections, baclofen injections, and the recovery period. Baclofen slightly increased the first wave amplitude but dramatically decreased the late wave amplitude ($n = 5$ animals, Friedman test with a *post-hoc* Dunn-Sidak test). Each data point indicates the average amplitude for a NeuroGrid electrode.

(D-F) We optogenetically inhibited wMC with NpHR during whisker touch while recording wS1 LFP with the NeuroGrids.

(E) (Top) Representative wMC spike raster during optogenetic inhibition. (Bottom) Average firing rates across 23 wMC single units during inhibition. These measurements were conducted in the absence of whisker touch.

(F) Peak-to-trough amplitude of the late wave across all electrodes during whisker touch with and without wMC optogenetic inhibition ($n = 3$ animals, $p < 0.001$, signed-rank Wilcoxon test). Each data point corresponds to a NeuroGrid electrode.

(G-H) We optogenetically stimulated wMC with ChR2 while recording the effects of the stimulation in wS1 with a silicon probe.

(H) Average wS1 LFP and corresponding CSD profile during optogenetic stimulation (left) and whisker touch (right) in the same animal. wMC optogenetic activation drives a delayed L2/3 current sink at ~200 ms, much in the same way that whisker touch does.
